# ILF2 blockade suppresses helicase-mediated R-loop resolution to induce lethality in homologous recombination-deficient cancers

**DOI:** 10.1101/2025.05.21.655390

**Authors:** Yi Chieh Lim, Soak Kuan Lai, Ru Han Yang, Yong Kang Chua, Rémi Patouret, Vaishnervi Manoj, Colin Chun Xuan Soh, Benjamin Lebeau, Ning Cham, Wei Meng, Sze Keong Tey, Szu Ling Yeap, Songjing Zhang, Jessie Yong Xing Woon, Zahin Ali, Marek Mutwil, Pritpal Singh, Melissa Jane Fullwood, Linda LD Zhong, Xue Ming Dong, Chiba Shunsuke, Cheng Gee Koh, Li Hoi Yeung

## Abstract

Hidden beneath the layer of tumor malignancy lies unchecked proliferation driven by hyperactive transcription, which fosters the formation of RNA:DNA hybrids. These structural intermediates precipitate collisions between the transcription and replication machineries. Central to mitigating this genomic threat is interleukin enhancer-binding factor 2 (ILF2), which orchestrates the recruitment of RNA:DNA helicases to resolve R-loops and, in doing so, reveals cancer cells dependency on stress-mitigating mechanisms to avert genome catastrophe. Our discovery of Molephantin and its lead derivative, NYH0002 is the outcome of delineating ILF2’s function. NYH002 exerts direct binding to the ILF2 complex and disrupts RNA:DNA helicase activity to elicit genome-wide DNA breakages. Tumors with elevated cyclin E and E2F1 expression are sensitive to NYH002, while those deficient in homologous recombination repair exhibit lethality. Collectively, these findings position ILF2 as a key regulator of R-loop homeostasis and reveal a therapeutic vulnerability that NYH002 exploits to suppress tumor malignancy.

## INTRODUCTION

There is an elegant symmetry between transcription and replication, where one decodes the genome while the other duplicates the DNA (1). Both processes are choreographed with remarkable spatial separation to ensure a harmonious interplay that does not compromise genome integrity. Oncogenic signaling on the other hand drives cancer cells into a state of hyper-transcription, increasing nascent RNA production to fuel uncontrolled proliferation at the consequence of fostering R-loops formation (2). These RNA:DNA hybrids are transiently formed within the active site of RNA polymerase when nascent RNA anneals to the template DNA strand and displaces the non-template strand, thus creating a three-stranded structure that is readily resolved upon the release of RNA (3). R-loops when left unregulated can obstruct advancing replication forks particularly in regions of the chromatin where transcription and replication machineries frequently converge, leading to transcription-replication conflicts (TRCs) (4, 5). Collisions between these two elements provoke replication stress, establishing an endless cycle of DNA damage-induced mutagenesis and tumor evolution (6). Paradoxically, excessive presence of R-loops pushes cancer cells toward a precarious state of overwhelming DNA damage, forcing them to rely on an overburdened post-replication repair system such as homologous recombination (HR) for survival (7, 8). The reliance on pathological R-loops to provoke replication stress-induced mutation and accelerate malignant progression is an exploitable vulnerability (9).

Therapeutic strategies that augment R-loop accumulation by disrupting their resolution can potentiate replication stress-induced DNA breakage through head-on TRCs. Although compounds such as romidepsin (HDAC inhibitor) (10) and camptothecin (TOP1 inhibitor) (11) have been proposed to enhance R-loop formation, they were not originally developed to target R-loop biology (12, 13). Rather, romidepsin and camptothecin ability to induce RNA:DNA hybrids arise as an indirect consequence of their primary mechanism of action. As of now, there are no clinically approved R-loop inducing drugs. Regardless of the strategies being employed, the deliberate objective is to surpass the DNA damage threshold essential for genomic stability and ultimately forcing a state of replication catastrophe. Persistent genomic stress can transform a tumorigenic mechanism into a lethal weakness (14). The resulting unresolved DNA DSBs offer a strategic basis for the selective elimination of cancer cells while sparing normal tissues with inherently low replication stress.

Here, we uncover a previously uncharacterized role of interleukin enhancer binding factor 2 (ILF2), which functions in concert with different helicases to resolve R-loop formation. Targeted inhibition of this pathway by NYH002, a derivatized Molephantin, induces selective cytotoxicity in cancer cells exhibiting G_1_/S-phase dysregulation, especially those with a compromised HR repair. Our discovery reveals insight into both the role of the ILF2 complex in preserving genomic stability and a therapeutic paradigm that exploits vulnerabilities in cancer-specific stress mitigation pathways to enhance targeted therapies.

## RESULTS

### Naturally Occurring Molephantin Displays High Therapeutic Potency and Cancer Selectivity

Harnessing cancer-specific toxicity for effective tumor eradication is a cornerstone of targeted therapy (15, 16) and serves as the underlying rationale for our retrospective observational study evaluating patient responses to integrative medicine involving orally administered phytotherapeutic agents (PAs). Of the 27 subjects evaluated, 20 were excluded due to ineligibility, withdrawal of consent, or non-compliance with the follow-up protocol (Figure S1A). Consequently, the cohort comprised three treatment-naïve patients and four post-first-line treated patients, all of whom had completed a minimum of a six-month integrative medicine regimen, with progression-free survival and disease regression evaluated as the primary and secondary outcomes, respectively. Over the 1.5-year follow-up period, routine clinical evaluations documented therapeutic benefits in five patients, with no observed adverse events (Figure S1B–C and 1A). Notably, three treatment-naïve patients exhibited a marked reduction in prostate-specific antigen (PSA) level and tumor size during the six-month treatment with the remaining patients demonstrating progression-free survival (Figure 1B–C).

**Figure 1:**
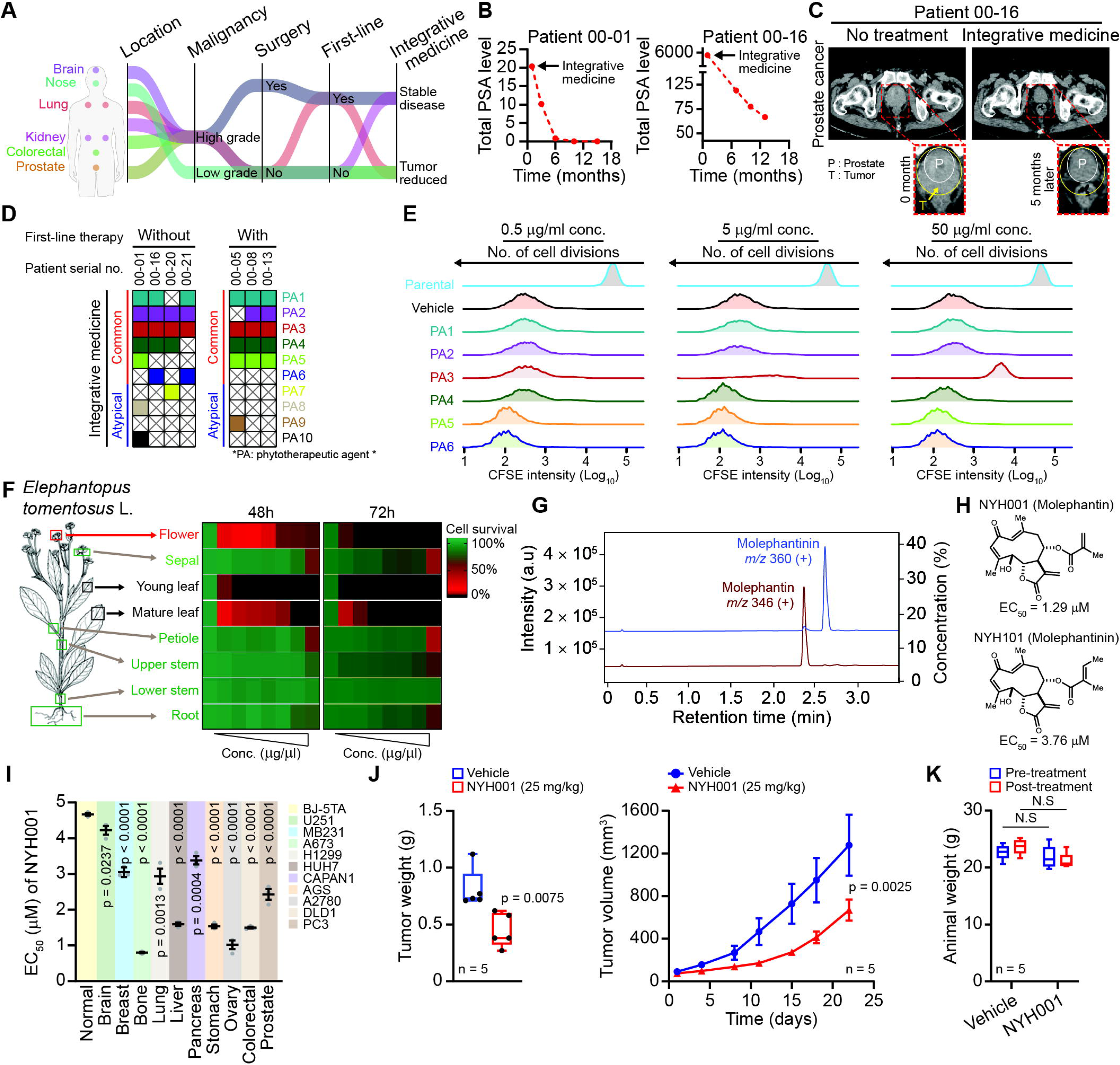
Identification of Molephantin revealed selective cytotoxicity in solid cancers. (A) Sankey plot of recruited cancer patients (n = 7) in a retrospective observational study on integrative medicine. (B) Progression of PSA level in naïve stage IV metastatic prostate cancer patients receiving integrative medicine. (C) CT scan of a naïve prostate cancer patient before and five months after receiving integrative medicine. ‘P’ denotes the region of the prostate, and ‘T’ indicates the region of the tumor. (D) Overall view of individual patients receiving integrative medicine, which comprised a minimum daily oral administration of combined PAs for a minimum period of six months. (E) Cell generation was tracked by measuring the fluorescence intensity of CFSE-labelled DLD1 cells treated with vehicle (DMSO) or 0.5, 5, and 50Jμg/ml of various PAs for 96 hours. “Parental” CFSE-labelled cells at day 0 were used as the reference for maximum fluorescence intensity in flow cytometry analysis (n = 3). (F) A simplified schematic of extract production was generated using various parts of *Elephantopus tomentosus* L., as indicated (left). Cell proliferation was determined at 48 and 72 hours after DLD1 cells received increasing concentration (0 – 1.70 mg/ml) of extract treatment (n = 3). (G) Isolation of Molephantin and Molephantinin compounds from young leaf extracts by preparative HPLC. (H) Chemical structures of Molephantin (NYH001) and Molephantinin (NYH101), and their dose-response effects in DLD1 cells. EC_50_ values were determined 48 hours after treatment (n = 3). (I) Cancer-specific cytotoxicity as revealed in a panel of tumorigenic (n = 10) and non-cancerous (n = 1) cell lines treated with a dose range of 0.1 – 10JμM of NYH001 for 48 hours. Data were presented as the average EC_50_ (n = 3). (J, K) Nude mice were engrafted with 1 × 10^6^ HCT116 cells, followed by intraperitoneal treatment with either vehicle or 25 mg/kg of NYH001. Treatment was performed every other day for two weeks with the animals’ weight measured daily. A control cull was executed upon completion of treatment, and individual tumors were excised for assessment. Data are presented as mean ± SEM. Unpaired two-tailed t-test (I, J, and K). p-values of < 0.05 were considered significant. See also Figure S1

Therapeutic activity based on the commonly administered PAs was subsequently evaluated in DLD1 cells where PA3 (*Elephantopus tomentosus L*.), demonstrated anti-proliferative properties (Figure 1D– 1F). This series of assays indicated growth inhibition, prompting the isolation of bioactive compound(s) from the crude extract. Sequential fractionation by flash column chromatography and monitored by thin-layer chromatography (TLC), revealed presence of distinct compounds across the collected fractions. Fractions displaying similar TLC profiles were consolidated and further refined by gel permeation chromatography. Subsequent nuclear magnetic resonance (NMR) spectroscopy identified the high purity and structural integrity of Molephantin and Molephantinin (Figure S1D and 1G). Designated as NYH001 (Molephantin) and NYH101 (Molephantinin), these natural germacranolide sesquiterpenes compounds exhibited effective therapeutic activity within the micromolar range. Between the two bioactive compounds, NYH001 demonstrated greater potency as a small-molecule inhibitor (Figure S1E and 1H), leading to the evaluation of its cancer-specific toxicity using a panel of 10 solid cancer cell-lines. BJ-5TA, a non-cancerous fibroblast cell line, was also included as a control in the assessment of therapeutic efficacy. The latter exhibited the least sensitivity to NYH001 (EC_50_ = 4.67 μM) as compared to the cancer cell line panel which responded within an EC_50_ range of 0.82 μM to 4.22 μM (Figure 1I). Furthermore, NYH001 treatment at 25 mg/kg in HCT116 tumor-bearing animal model demonstrated consistent therapeutic efficacy in mitigating tumor progression while exerting minimal systemic cytotoxicity (Figure 1J–K).

### NYH002 Suppresses Cancer Growth by Disrupting DNA Integrity

The isolated NYH001 molecule possesses a rare 10-membered macrocyclic backbone that includes an (E,Z)-dienone moiety spanning carbon atoms 1 to 4. Crucially, this ring structure is fused to a lactone ring known as an α-methylene-γ-butyrolactone, and the molecule encompasses a cluster of four contiguous stereogenic centers at carbon atoms 5 to 8 (17). This intricate chemical structure suggests the molecule may have unique biological function, as demonstrated by its earlier observed ability to selectively induce cytotoxicity in solid cancer cell lines while sparing non-cancerous cells. To investigate the molecular mechanism of Molephantin, we employed a structure–activity relationship (SAR) approach, leveraging the chemical versatility of aromatic carboxylate scaffolds to first develop lead compounds. Five different derivatives, designated NYH002 – NYH006, were synthesized with structural variations to improve interactions with nucleophilic residues within the target protein, thereby enhancing therapeutic efficacy (Figure S2A–B and 2A). The design of our strategy enhances the drug-protein binding affinity while simultaneously reducing off-target interactions.

Treatment of a panel of 10 solid cancer cell lines (U251, MB231, A2780, H1299, HUH7, CAPAN1, AGS, A673, DLD1 and PC3) with NYH002 – NYH006, revealed these compounds could achieve an EC_50_ at nanomolar range (Figure 2B). All five derivatives exhibited improved efficacy with NYH002 having the greatest potency (Figure 2C). For a therapeutic safety evaluation, animals were daily administered with the top two lead candidates, NYH002 and NYH003 by intraperitoneal route for two weeks (Figure 2D). Not only did NYH002 show favorable tolerability *in vivo*, its therapeutic efficacy to elicit cancer cell death also exceeded that of the original NYH001 (Figure 2E).

**Figure 2:**
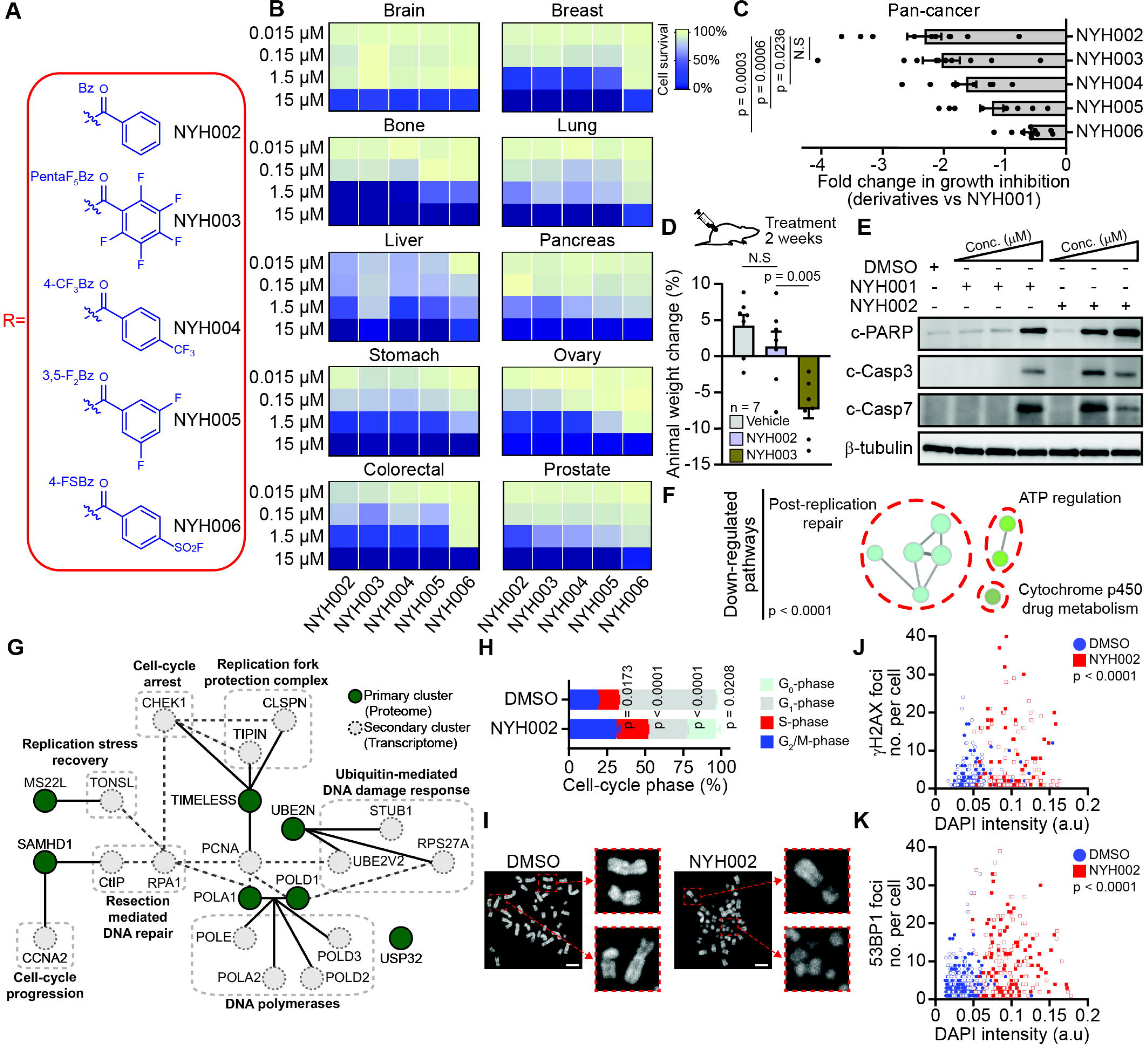
Lead-optimized NYH002 suppresses replication machinery and causes chromosomal aberrations. (A) Structural analogue of NYH001 derivatives. (B) Dose-response of NYH001 derivatives (NYH002 – NYH006) in a panel of 10 different cancer cell lines as indicated. Mean survival was determined at 48 hours post-treatment (n = 3). (C) Therapeutic efficacy of NYH001 derivatives. Fold change in mean EC_50_ was assessed across 10 cell lines and presented as pan-cancer data at 48 hours post-treatment. Each derivative was compared to the corresponding NYH001-treated cell line, where individual dot represents the mean EC_50_ of the respective cancer cell line (n = 3). (D) Evaluation of systemic cytotoxicity. A total of 21 healthy nude mice was randomly assigned to vehicle, NYH002 and NYH003 treatment arms. Each animal received a total of 14 daily intraperitoneal injections at 25 mg/kg for 2 weeks. Animal weights were recorded before and after treatment to assess animal health. (E) Drug-induced cell death in DLD1 cells following treatment with increasing concentrations of NYH001 and NYH002 (1 – 10JμM) for 24 hours. An immunoblot was performed with WCL to determine activation of apoptosis by cleavage of PARP, caspase-3, and caspase-7. β-tubulin was used as a loading control (F) Pathway enrichment analysis of the downregulated proteome was performed on DLD1 cells treated with 2.5JμM NYH002 for 24 hours, followed by TMT-labeling of WCL for MS analysis (n = 3). Based on KEGG, Reactome, WikiPathways, and Gene Ontology databases, the differentially downregulated proteins were subjected to pathway enrichment analysis. (G) The protein–protein interaction (PPI) network of NYH002-treated DLD1 cells was established using the STRING and IntAct databases. The network was built on the downregulated proteome as the first interactors (primary clusters), followed by the integration of downregulated transcriptome data to identify the second interactors (secondary clusters). (H) Cell cycle profile based on the propidium iodide staining of DMSO- and 2.5JμM NYH002-treated DLD1 cells after 24 hours (n = 3). (I) Chromatid breaks in metaphase spread of DLD1 cells treated with DMSO or 2.5JμM NYH002 for 24 hours (n = 3). Scale bar: 5 μm. (J, K) Quantitative image-based cytometry of immunolabeled DLD1 cells receiving either DMSO or 2.5JμM NYH002 for 24 hours. Total number of γH2AX or 53BP1 foci and nuclear DAPI intensity were determined for each of >1,000 individual cells (n = 3). Data are presented as mean ± SEM. Unpaired two-tailed t-test (C, D and H), hypergeometric test with Bonferroni step-down (F) and Mann-Whitney U test (J and K). p-values of < 0.05 were considered significant whereas N.S was regarded as not significant. See also Figure S2

We next utilized a system biology approach to determine the signaling network perturbed by NYH002. DLD1 cells received NYH002 treatment followed by tandem mass tag (TMT) labeling in conjunction with liquid chromatography-tandem mass spectrometry (LC-MS/MS) analysis. A total of 6,297 labeled proteins were then filtered for differentially expressed proteins of which 71 down-regulated proteins were determined (Figure S2C–D). Pathway enrichment analysis of this downregulated protein set indicated NYH002 as a disruptor of post-replication repair (Figure S2E and 2F). To substantiate this observation, transcriptome analysis was also conducted to complement the results of the pharmacoproteomics assessment (Figure S2F). Integration of both omics datasets supports a potential role for NYH002 in modulating the replication machinery (Figure 2G). This prompted the investigation into the integrity of DNA synthesis. Following 24 hours of NYH002 treatment, DLD1 cells exhibited G_2_/M phase arrest, a critical checkpoint for resolving DNA strand breakages prior to cell division (18). Notably, NYH002-treated DLD1 cells exhibited gross chromosomal breaks resulting from unresolved DNA lesions (Figure 2H–I). Supporting this observation, we detected the presence of γH2AX and 53BP1 foci, which are well-established markers of DNA DSBs and are commonly recruited to sites of stalled replication forks (19, 20). In NYH002-treated DLD1 cells, γH2AX and 53BP1 foci localized within intensely DAPI-stained nuclei (Figure 2J–K), suggesting a direct link to replication stress rather than oxidative stress-induced DNA damage (Figure S2G).

### DNA Strand Breaks Arise from Collisions Between Transcription and Replication Machinery

Building on our observed evidence of DNA DSBs occurring in a cell cycle-dependent manner, we systematically investigated whether NYH002 induces replication stress. Pulse-chase imaging using our NYH002-alkyne (NYH002P) azide-activated fluorescent (AF594) revealed rapid nuclear localization of the compound with minimal detection in the cytoplasm (Figure S3A and 3A). Synchronized G_1_/S-phase DLD1 cells, upon release from aphidicolin (APH) and treatment with NYH002, exhibited high accumulation of γH2AX foci when progressing into S-phase (Figure S3B–E and 3B). In contrast, CDK1 (RO-3306) inhibitor synchronized G_2_/M-phase DLD1 cells which were subjected to the same treatment condition and transitioning into G_1_-phase showed negligible DNA strand breakages. Furthermore, DNA fiber combing demonstrated a lack of ongoing replication and increased fork stalling (Figure S3F and 3C). This series of findings provides compelling evidence that NYH002 acts in proximity to the replication machinery to facilitate fork collapse.

We therefore conducted high-throughput chromosome conformation capture (Hi-C) sequencing on NYH002-treated DLD1 cells to examine structural abnormalities in chromatin organization to delineate the mechanisms underlying replication dynamics disruption. In each of the depicted gene sets, alterations in topologically associating domains (TADs) at their transcription start sites (TSSs) remained unchanged, except for the downregulated gene set (Lost <1), which exhibited a gain in chromatin interactions (Figure 3D). The high-contact-frequency chromatin compaction is restricted to short genomic distances, such as those identified in *NFKB1* locus where there was a shift in loop interaction (Figure 3E–F). These condensed nuclear regions, marked by H3K27me3 expression (Figure 3G), could be a direct consequence of RNA polymerase II (RNAPII) stalling caused by TRCs (21). To this end, immunoblotting of NYH002-treated DLD1 cells confirmed the retention of RNAPII, which also coincided with the induction of DNA DSBs and chromatin condensation, as evidenced by γH2AX activation and increased HP1 expression, respectively (Figure 3H).

**Figure 3:**
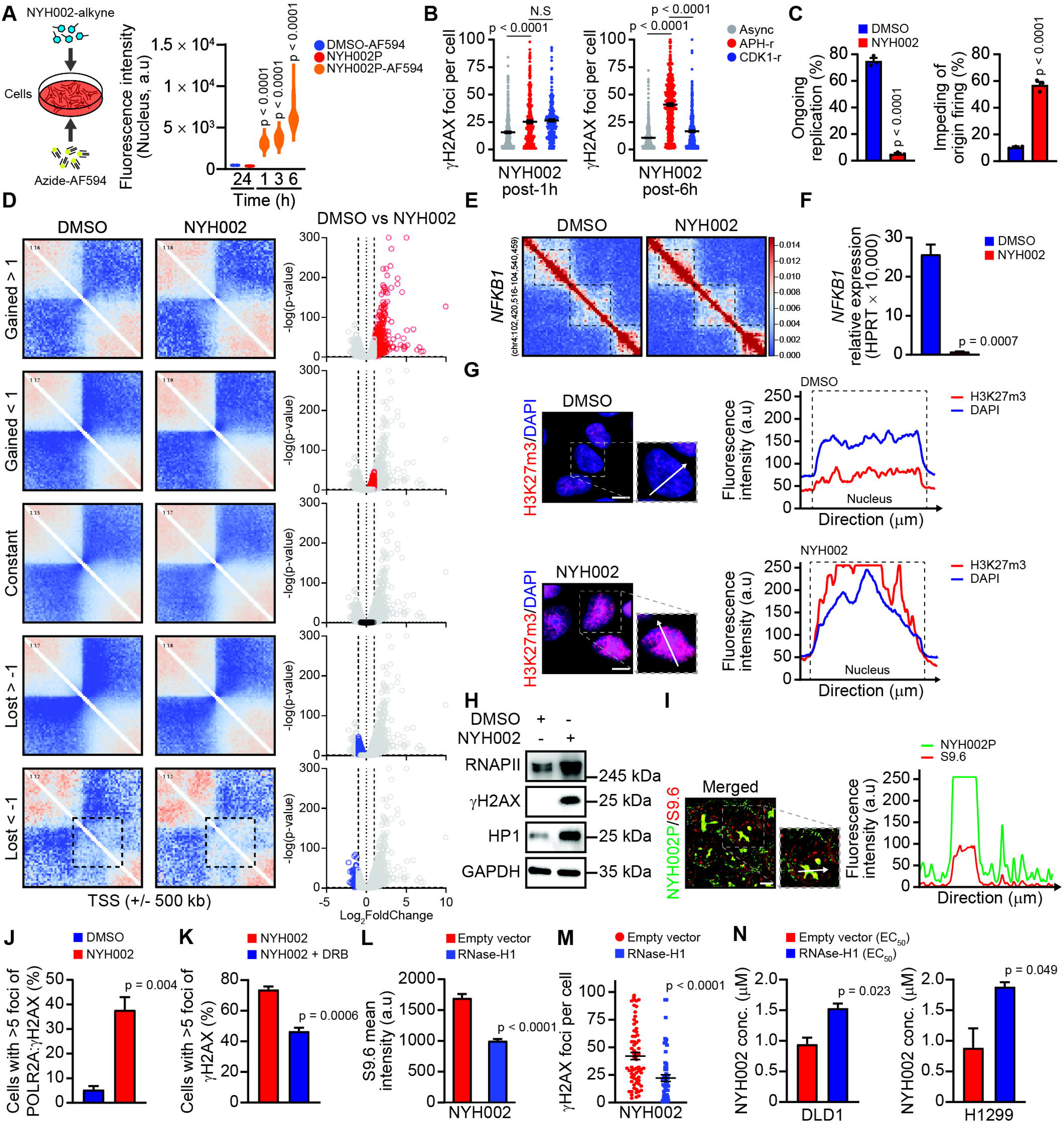
NYH002 induces replication breakages via the collision of pathological R-loops. (A) A simplified schematic strategy to determine NYH002 localization in the cellular compartment (left). NYH002-alkyne (NYH002P) was enabled by click chemistry to function as a fluorescent probe. DLD1 cells were treated with 0.5JμM of this compound and fixed at the indicated time points to determine NYH002P localization within the nuclear compartment. (B) Replication-induced fork breakage was assessed by synchronizing DLD1 cell at G_1_/S-phase with APH or G_2_-phase with CDK1 inhibitor for 16 hours. Synchronized cells were released from these chemicals (APH-r and CDK1-r) and immediately received 2.5JμM of NYH002 treatment. DLD1 cells were fixed at one- and six-hours post-treatment and analyzed for DNA DSB by γH2AX foci count. (C) Replication fork stalling was assessed by quantifying the total number of ongoing DNA fiber (left) events and the lack of origin firing (right) in DLD1 cells treated with either DMSO or 2.5JμM NYH002 (n = 3). (D) Pile-up interaction at the TSS of genes with altered or constant expression following NYH002 treatment (left), alongside a volcano plot showing RNA expression of the genes represented within the pile-up (right). The black dashed square indicates a region of increased interaction at the TSS of downregulated genes following 2.5JμM NYH002 treatment. (E) Hi-C heatmap at 32kb resolution centered around the downregulated *NFKB1* gene by NYH002 treatment showing distinct effect of chromatin condensation. (F) Transcript expression of *NFKB1* gene in DLD1 cells was determined by RT-qPCR after 24 hours treatment with DMSO or 1JμM of NYH002 (n = 3). (G) Chromatin condensation was determined by H3K27m3 expression in the nucleus of DLD1 cells treated with DMSO or 1JμM of NYH002 for 24 hours (n = 3). Arrow in white indicates orientation of direction. Scale bar: 5 μm. (H) Immunoblot of transcription stalling in DLD1 cells treated with DMSO or 1JμM NYH002 for 24 hours, as determined by RNAPII, γH2AX and HP1 expression levels. GAPDH was used as loading control. (I) Presence of R-loops and their proximity to NYH002 localization were assessed 24 hours after treatment with 1JμM of the compound’s probe, followed by click chemistry with AF549 fluorophore and immunofluorescence staining using the S9.6 antibody in DLD1 cells. Arrow in white indicates orientation of direction. Scale bar: 5 μm. (J) Collision between replication and transcription machinery was determined by the co-localization events of RNAPII and γH2AX foci post-24 hours after treatment with DMSO or 1JμM NYH002 in DLD1 cells (n = 3). (K) Dissolution of TRC through transcription inhibition by DRB (50JμM) was carried out one hour prior to the treatment of 1JμM NYH002 in DLD1 cells. Cells were examined 24 hours later for the presence of DNA DSBs by quantifying total cell number with >5 γH2AX foci (n = 3). (L, M) Phenotypic profile of R-loops was assessed in DLD1 cells transfected with empty vector (EV), or RNaseH1 for 24 hours, followed by treatment with 1JμM NYH002 for an additional 24 hours. Immunofluorescence staining was then performed using either the S9.6 or γH2AX antibody. (N) Reduction in NYH002 efficacy was assessed in a dose-dependent manner in DLD1 and H1299 cells transfected with either RNaseH1 to resolve R-loops or with an empty vector. Cell viability was measured 48 hours post-treatment and data were presented as mean EC_50_ of NYH002 (n = 3). Data are presented as mean ± SEM. Mann-Whitney U test (A, L and M). Unpaired two-tailed t-test (B, C, J, K and N). p-values < 0.05 were considered significant whereas N.S was regarded as not significant. See also Figure S3

NYH002 likely triggers TRCs through its ability to promote the formation of RNA:DNA hybrid structures (Figure S3G–H and 3I) (22). To test this hypothesis, we examined the spatial proximity between the transcription machinery and replication fork collapse by assessing the co-localization of RNAPII and γH2AX foci, respectively in NYH002-treated DLD1 cells (Figure 3J) (22). In line with this approach, transcription inhibition using DRB significantly reduced replication-dependent DSBs, which is consistent with TRC-driven DNA damage in response to NYH002 treatment (Figure 3K). The TRC-dependent DSB-induced genome instability raised the question of whether R-loop removal would disrupt NYH002’s mechanism of action. Hence, DLD1 and H1299 cells were transduced with a RNaseH1-expressing vector to catalyze the cleavage of ribonucleotides in R-loop substrates, followed by NYH002 treatment. We observed that RNaseH1-expressing cancer cells exhibited a marked reduction in γH2AX foci accumulation after NYH002 treatment, which was accompanied by improved overall survival, as revealed by the higher EC_50_ of NYH002 (Figure S3I and 3L–N).

### NYH002 Targets R-Loop Dynamics by Modulating ILF2-Dependent Helicase Function

By disrupting R-loop resolution, NYH002 promotes unresolved pathological RNA-DNA hybrids that directly obstruct replication fork progression. Our findings indicate NYH002 targets a core resolving factor critical to maintaining R-loop homeostasis. Hence, we conducted a competitive binding assay pairing NYH002 (antagonist) with its biotinylated derivative (NYH002P, interactor) to identify this target of interest (Figure S4A and 4A). Our data revealed a single high-affinity protein interactor between 48 kDa and 35 kDa in molecular weight (Figure 4B). This unidentified high-affinity (core) interactor of NYH002P, along with the remaining proteins (associate interactors), was subjected to LC-MS/MS analysis to determine the composition of the protein complex. (Figure S4B). Peptide identification revealed a total of 795 interacting proteins. However, only 183 of these interacting proteins corresponded to the expected molecular weights of the excised silver-stained gel band regions (Figure S4C and 4C). We then leveraged the core and associated interactors to sequentially map protein-protein interactions (PPIs) based on reference nuclear complexes from the CORUM database (23). The annotation strategy revealed a distinct interaction network enriched in DNA/RNA-binding proteins and transcription regulators (Figure S4D and 4D).

**Figure 4:**
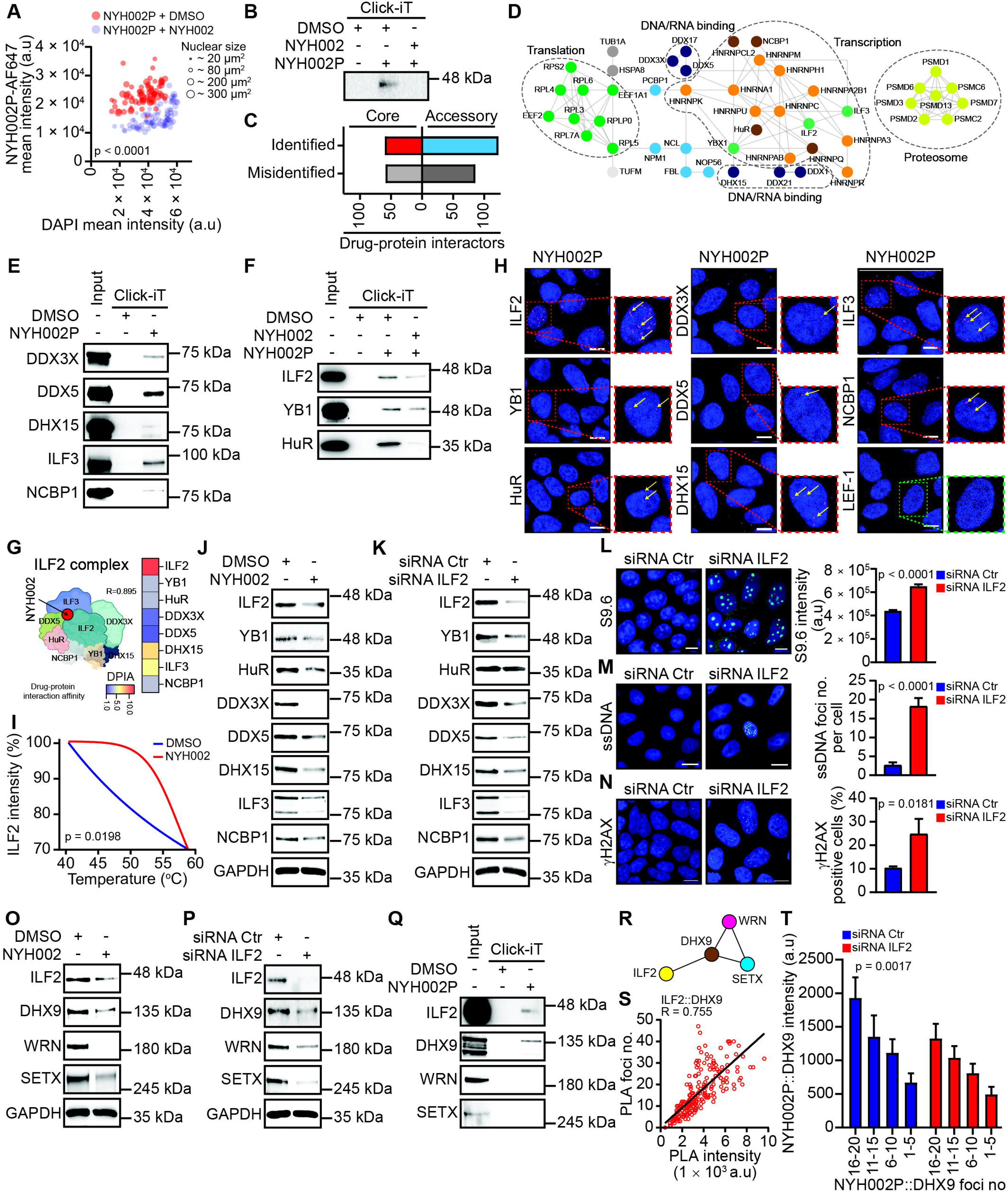
Targeting the ILF2-DHX9 axis by NYH002 increases R-loop accumulation. (A, B) Competitive binding involved the initial treatment with NYH002 (2JμM) or DMSO for one hour, followed by the addition of NYH002P (1JμM) for an additional five hours. Click chemistry was performed between NYH002P and the AF647 fluorescent probe for immunofluorescence analysis in DLD1 cells, or alternatively with biotin for subsequent HRP-based detection in WCL by immunoblotting. (C) MS label-free quantification of gel-separated proteins was performed on NYH002P-biotin co-immunoprecipitation from DLD1 WCL. Direct (core) and indirect (accessory) drug–protein interactors were classified as either ‘identified’, which matched the expected molecular weight, or ‘misidentified’, which showed an incorrect molecular weight (n = 3). (D) NYH002P–protein interactome was curated using the nuclear complex dataset from CORUM followed by the construction of a PPI network with STRING. (E) Validation of NYH002-interacting proteins that were identified from the ‘DNA/RNA binding’ and ‘Transcription’ cluster in STRING. Co-immunoprecipitation was performed using click chemistry between NYH002P and biotin in DLD1 cells. Input represents 1% of the total WCL. (F) Competitive binding of NYH002P-interacting proteins was assessed by co-immunoprecipitation in DLD1 cells. Input represents 1% of the total WCL. (G) A simplified model based on AlphaFold3 prediction of the ILF2 protein complex. (H) Visualization of drug–protein interactions. PLA was carried out between the indicated proteins and post-treatment with NYH002P (1JμM) at six hours in DLD1 cells. Foci of PLA in the nucleus represent physical drug–protein interaction. LEF-1 nuclear protein was used as a negative control (n = 3). Scale bar: 10 μm (I) Cellular thermal shift assay was employed to determine the direct interaction between NYH002 and ILF2. DMSO was used as a negative control (n = 3). (J, K) Immunoblot of the ILF2 complex revealed that treatment with 1JμM NYH002 for 24 hours in DLD1 cells phenocopied the effect of ILF2 depletion by siRNA. (L, M, N) Phenotypic characterization of R-loops in siRNA-mediated ILF2-depleted DLD1 cells at 48 hours post-transfection. Loss of ILF2 resulted in the induction of S9.6, ssDNA, and γH2AX foci (n = 3). Scale bar: 10 μm. (O, P) Immunoblot showing downregulation of DHX9, WRN, and SETX helicases as a direct consequence of ILF2 targeting by NYH002 (1JμM, 24 hours post-treatment) or by siRNA-mediated depletion (48 hours post-transfection). GAPDH was used as a loading control. (Q) Co-immunoprecipitation of NYH002P-biotin via click chemistry revealed a direct interaction with ILF2 and DHX9, but neither with WRN nor SETX. Input represents 1% of the total WCL derived from DLD1 cells. (R) *In silico* STRING analysis indicated that the interaction of ILF2 with WRN and SETX is dependent on DHX9. (S) PLA in the nucleus of DLD1 cells showed direct interaction between ILF2 and DHX9 (n = 3). (T) siRNA-mediated ILF2 depletion in DLD1 cells resulted in a significant reduction in PLA foci formation between NYH002P and DHX9 (n = 3). Data are presented as mean ± SEM. Mann-Whitney U test (A, L and M), two-way ANOVA (I and T) and unpaired two-tailed t-test (N). p-values < 0.05 were considered significant. See also Figure S4

Immunoprecipitation of NYH002P utilizing whole cell lysate (WCL) was conducted to investigate all uniquely identified DNA/RNA-binding and transcription-related proteins, including one representative each from the translation cluster and heterogeneous nuclear ribonucleoproteins. The analysis uncovered a novel protein complex composed of eight distinct members (Figure S4E and 4E–F). Independent assessment based on amino acid sequence among the eight candidate proteins confirmed a high confidence interacting complex (Figure S4F and 4G). Notably, the molecular weights of ILF2, YB1, and HuR were consistent with the earlier unidentified immunoblot band range (<48 kDa and >35 kDa), positioning these proteins as plausible core interactors of NYH002. In a follow-up assessment using the proximity ligation assay (PLA) to establish spatial proximity within the uncharacterized complex, ILF2 emerged as the primary interacting partner of NYH002P (Figure S4G– H and 4H). The cellular thermal shift assay (CESTA), which evaluates protein thermostability in the presence of drug binding, provided further confirmation of the physical interaction between ILF2 and NY002 (Figure 4I) (24).

Accordingly, we investigated the mechanism NYH002 modulates ILF2 activity to disrupt the stability and assembly of the protein complex. Immunoblot analysis revealed that the treatment with NYH002 did result in a coordinated downregulation of all members of the complex, a pattern that was recapitulated upon ILF2 knockdown via siRNA (Figure 4J–K). Notably, ILF2 depletion in DLD1 cells alone was sufficient to recapitulate phenotypic features of NYH002’s mechanism, namely the accumulation of unresolved R-loops and the displacement of single-stranded (ss)DNA (Figure 4L–M). This series of phenotypic observations supports the role of ILF2 as a mediator of replication stress which ultimately led to fork collapse as evidenced by the accumulation of γH2AX foci (Figure 4N).

Despite ILF2 having been previously suggested in R-loop regulation, there were discrepancies regarding the molecular mechanism through which it acted (25, 26). Evidence supporting ILF2 as a central facilitator in coordinating RNA/DNA-binding proteins to regulate R-loop emerged from our earlier PLA data. Building on this mechanistic insight, we investigated key RNA:DNA helicases known to mediate R-loop resolution (27). Interestingly, immunoblot analysis of NYH002-treated and ILF2-depleted DLD1 cells revealed similar downregulation effect of DExH-box helicase 9 (DHX9), Senataxin (SETX), and Werner syndrome helicase (WRN), all major helicases that are essential for maintaining R-loop homeostasis (Figure 4O–P) (28, 29). However, immunoprecipitation of NYH002P showed direct interaction with DHX9 but not with SETX or WRN (Figure 4Q). PPI based on STRING analysis supported the argument that ILF2 regulates SETX and WRN through DHX9 interaction (Figure 4R). This prediction was corroborated by our immunoprecipitation and PLA results, which showed a direct interaction between DHX9 and both SETX and WRN (Figure S4I–K and 4S). In contrast, ILF2 did not exhibit physical binding to either SETX or WRN (Figure S4L–M). These findings therefore reinforce the notion that NYH002 impairs R-loop resolution by targeting ILF2 (Figure 4T).

### NYH002 Demonstrates Therapeutic Potential in Both Monotherapy and Combination Treatment

Elucidating NYH002’s mechanism of actions reveals its potential as a concomitant therapy to enhance first-line DNA-damaging treatments by enhancing irreversible DNA strand breakages to trigger apoptosis, despite presence of an intact post-replication DNA repair system (Figure S5A). To test this rationale, we leverage the enhancement of RNA:DNA hybrid accumulation to exacerbate R-loop burden while simultaneously introducing non-toxic DNA lesions to promote replication stress. NYH002 in combination with clinically-approved chemotherapeutic agents – temozolomide (TMZ) and 5-fluorouracil (5FU), with methyl methanesulfonate (MMS) serving as a positive control for replication stress, demonstrated synergistic effects (Figure 5A) (30, 31). At sub-therapeutic concentrations that do not elicit DNA damage-mediated apoptosis, chemotherapeutic agents (TMZ, 5-FU, and MMS) were still capable of inducing replication stress-induced DSBs (Figure 5B–C). Similarly, treatment with NYH002 at approximately its EC_25_, which was chosen to avoid triggering apoptosis in DLD1 cells, resulted in minimal induction of γH2AX foci. In contrast, the combination of NYH002 with either TMZ, 5-FU, or MMS led to a pronounced accumulation of replication stress-induced DSBs and apoptosis, as evidenced by the high degree of unresolved γH2AX foci and elevated levels of cleaved caspase-3, respectively. These findings highlight a synergistic interaction between NYH002 and DNA-damaging agents in amplifying the apoptotic response beyond what is observed with either therapeutic agent alone. As a proof-of-concept to validate the clinical relevance of this concomitant therapeutic strategy, we investigated the most effective combination, TMZ and NYH002, at 25 mg/kg and 10 mg/kg, respectively, in an animal model. Administration of alternate-day treatment over the initial two-week period resulted in complete inhibition of tumor progression. Consequently, the concomitant treatment regimen of TMZ and NYH002 was extended for an additional seven weeks to determine the possibility of therapeutic resistance, during which no palpable tumor masses were detected (Figure 5D).

**Figure 5:**
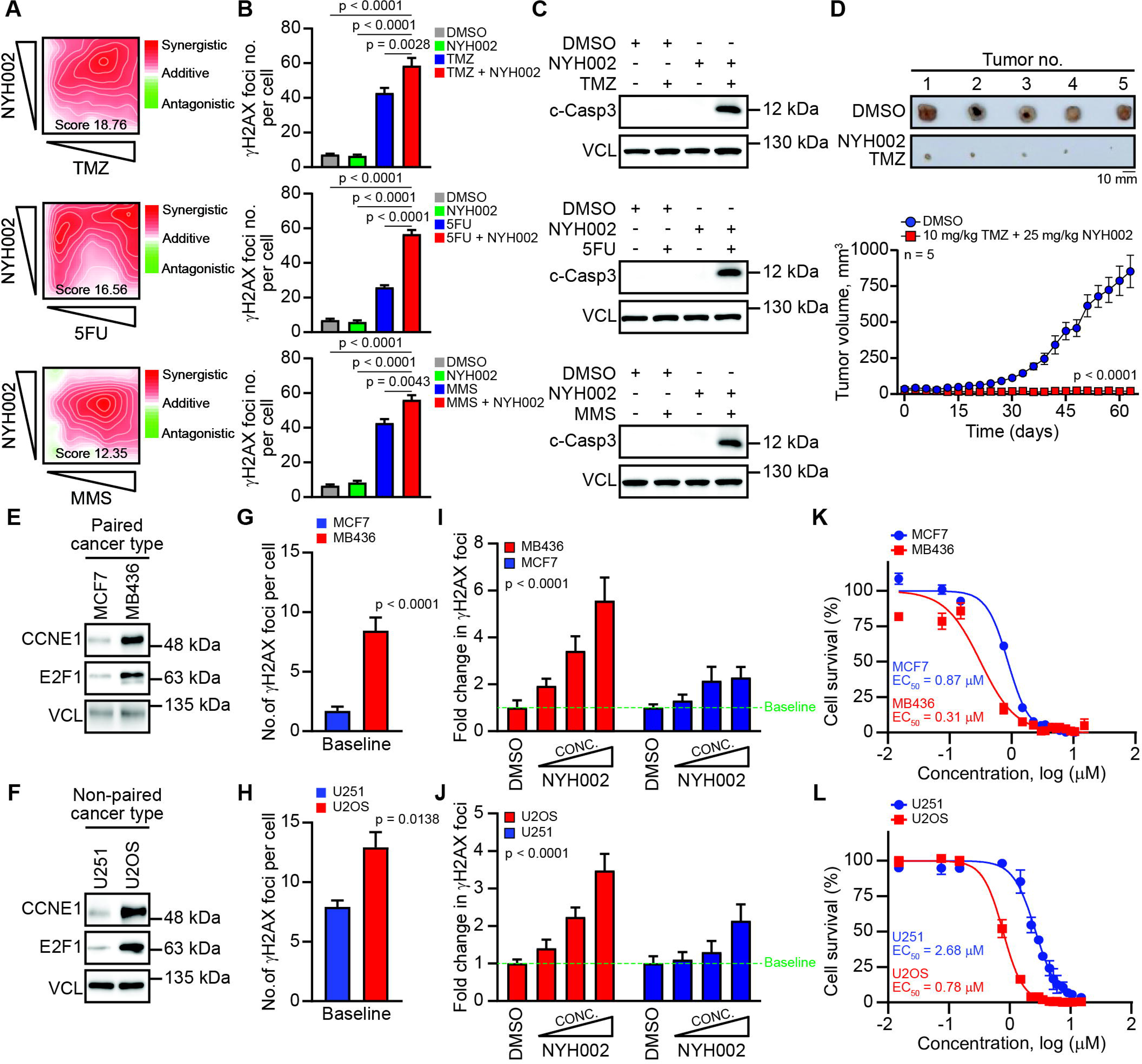
NYH002 synergizes with DNA-damaging agents to suppress tumor progression and exhibits monotherapy efficacy against cancer cells with replication stress. (A) A series of dose–response matrices of NYH002 in combination with TMZ, 5-FU, or MMS. Synergy map showed the combination effects of DLD1 cell survival as indicated by SynergyFinder 3.0. A score of >10 indicates synergistic, <10 indicates additive, and <-10 indicates antagonistic. (B, C) Synergistic effect of NYH002 with TMZ, 5-FU, or MMS on the induction of DNA DSBs and apoptosis. DLD1 cells received 0.5JμM NYH002, 1.5JmM TMZ, 0.1JmM 5-FU, or 0.5JmM MMS as monotherapies or in combination, as indicated, for 24 hours prior to immunostaining for γH2AX. Alternatively, WCL were prepared for immunoblotting to examine cleaved caspase-3. Vinculin was used as a loading control. (D) Synergy of NYH002 and TMZ *in vivo*. Flank tumor implantation was performed by injecting 1 x 10^6^ cells into the subcutaneous space of the animals, followed by intraperitoneal dosing of NYH002 in combination with oral administration of TMZ for nine weeks. Tumor growth was monitored and measured daily. A control cull was performed at day 63. (E, F) Immunoblot of differential protein expression of CCNE1 and E2F1 as indicators for G_1_/S checkpoint deficiency. MCF7 and MDA-MB436 cell lines represent a matched tumor pair for breast cancer, whereas U251 and U2OS serve as a non-paired tumor comprising of glioblastoma and osteosarcoma, respectively. Vinculin was used as a loading control. (G, H) Baseline replication stress is indicated by the presence of unresolved γH2AX as shown by the different cell lines (n = 3). (I, J) Increased susceptibility to replication-induced DSBs after treatment with 0.5 – 1.5 μM of NYH002 for 24 hours was observed in MDA-MB436 and U2OS cells with G_1_/S-phase checkpoint deficiency, as compared to MCF7 and U251 cells with normal cell cycle progression (n = 3). (K, L) EC_50_ of NYH002 was determined after 48 hours in both paired-match (MCF7 and MDA-MB436) and non-paired (U251 and U2OS) cell lines (n = 3). Data are presented as mean ± SEM. Mann-Whitney U test (B, G and H), two-way ANOVA (D) and one-way ANOVA (I and J). p-values < 0.05 were considered significant. See also Figure S5

Current evidence supporting this concomitant therapeutic approach reinforces the notion that G_1_/S checkpoint-deficient tumors, characterized by their intrinsic genetic susceptibility to replication-induced DSBs (32), are likely to exhibit heightened sensitivity to NYH002 monotherapy. The identification of predictive marker/s for G_1_/S checkpoint deficiency (Figure S5B–C), demonstrated a strong correlation among *CCNE1*, *E2F1*, *E2F2*, and *CDC25A* genes (Figure S5D) which included their significance in stratifying patient prognosis (Figure S5E). Leveraging on the identification of *CCNE1* and *E2F1* as top predictive markers for NYH002 monotherapy in G_1_/S checkpoint-deficient tumors as supported by their association with *RB1* mutation (Figure S5F), we curated the Cancer Cell Line Encyclopedia (CCLE) database to identify tumor models exhibiting differential expression of both genes. To establish a pan-cancer model system, we selected two pairs of cell-lines for comparative analysis. The first pair-match, MCF7 and MDA-MB-436 cells, represents breast cancer as a lineage-specific tumor model. The second non-pair match, comprising glioblastoma U251 and osteosarcoma U2OS cells, will serve as a comparison across tumor types (Figure S5G). Together, these four cancer cell lines will establish a therapeutic framework for targeting G_1_/S checkpoint deficiency in a pan-cancer context. Immunoblot analysis confirmed elevated expression levels of *CCNE1* and *E2F1* in MDA-MB-436 and U2OS cells (Figure 5E–F), which were consistent with a higher baseline of unresolved DNA damage associated with replication stress (Figure 5G–H). Upon treatment with increasing concentrations of NYH002, which targets R-loop resolution, both MDA-MB-436 and U2OS cells displayed sensitivity to the inhibitor, as indicated by the progressive accumulation of unresolved γH2AX foci (Figure 5I–J) and a positive S9.6 fluorescence signal (Figure S5H). Consequently, these checkpoint-deficient cancer cells, which were unable to efficiently resolve replication stress-induced DSBs, exhibited a significant reduction in overall survival (Figure 5K–L). In contrast, MCF7 and U251 cells, which possess a functional G_1_/S checkpoint, displayed reduced sensitivity to NYH002 under identical treatment conditions. In these cancer cells, γH2AX foci were efficiently resolved despite increasing concentrations of NYH002. Furthermore, MCF7 and U251 cells showed better treatment tolerance, as reflected by higher EC_50_ values, highlighting the vulnerability of G_1_/S checkpoint-deficient cells to NYH002 therapy.

### NYH002 Exploits DNA Repair Vulnerabilities to Unmask Therapeutic Potential in Homologous Recombination Deficient Cancers

The marked difference in therapeutic sensitivity of G_1_/S checkpoint-deficient cancer cells to NYH002, compared to their counterparts with functional cell cycle checkpoint, likely reflects a compromised DDR. Such impairment reduces cancer cells’ ability to repair replication stress-induced DSBs induced by NYH002 (33). By analyzing a curated cohort of 9,880 solid tumors in a pan-cancer setting, we established a positive correlation between high *CCNE1* and *E2F1* expression and the presence of 70 and 34 mutated DDR genes, respectively (Figure S6A–B). In contrast of this observation, differential expression of CCND1 and CDK4, which governed the G_1_-phase cell cycle progression did not correlate with DDR mutations (Figure S6C–D). Venn diagram analysis illustrated that 31 genes were common between high expressing *CCNE1* and *E2F1* (Figure S6E–I). Using this common DDR gene set, pathway enrichment concluded that most G_1_/S checkpoint-deficient tumors had a limited HR repair capacity (Figure S6J–K). Hence, we investigated the context of pathway lethality by performing a loss-of-function analysis using a siRNA library that targets the DDR cascade (n = 239). This independent yet unbiased approach enabled a systematic evaluation of the functional contributions of key genes within the DDR network. Our findings identified 209 knockdown genes that conferred selective vulnerability to NYH002-treated DLD1 cells, with the majority implicated in the response to DSBs (Figure 6A–B). Pathway enrichment using the top 90th percentile of DDR genes revealed a critical role of HR in cancer cell’s survival (Figure 6C). Recognizing the pivotal role of a limited post-replicative repair mechanism in mediating susceptibility to NYH002 treatment, we investigated the top three candidate HR genes, exonuclease 1 (*EXO1*), mutS homolog 3 (*MSH3*) and flap structure-specific endonuclease (*FEN1*) that were identified by the siRNA DDR library for synthetic lethal interaction (Figure S6L and 6D) (34–36). Here, we moved beyond the reliance on breast cancer (BRCA) 1/2 mutations and utilized genomic scar assessment (Figure S6M) to capture a broader spectrum of HR deficiencies (37). Using a cut-off score of ≤42, we interrogated the CCLE database to identify representative cancer cell lines from each major solid tumor type exhibiting HR proficiency (38, 39). An HR-deficient (HRD) cell line (score >42) was included as a negative control (Figure S6N). Notably, siRNA-mediated depletion of *EXO1*, *MSH3*, or *FEN1* exacerbated NYH002 sensitivity (approximately EC_25_) in all HR-proficient cancer cell lines, resulting in near-complete ablation of cell growth (Figure 6E). This effect was, however, absent in the HRD CAPAN-1 cell line, which had already exhibited pre-existing susceptibility to NYH002. Further depletion of *EXO1*, *MSH3*, or *FEN1* did not exacerbate growth inhibition in NYH002-treated CAPAN-1 cells.

**Figure 6:**
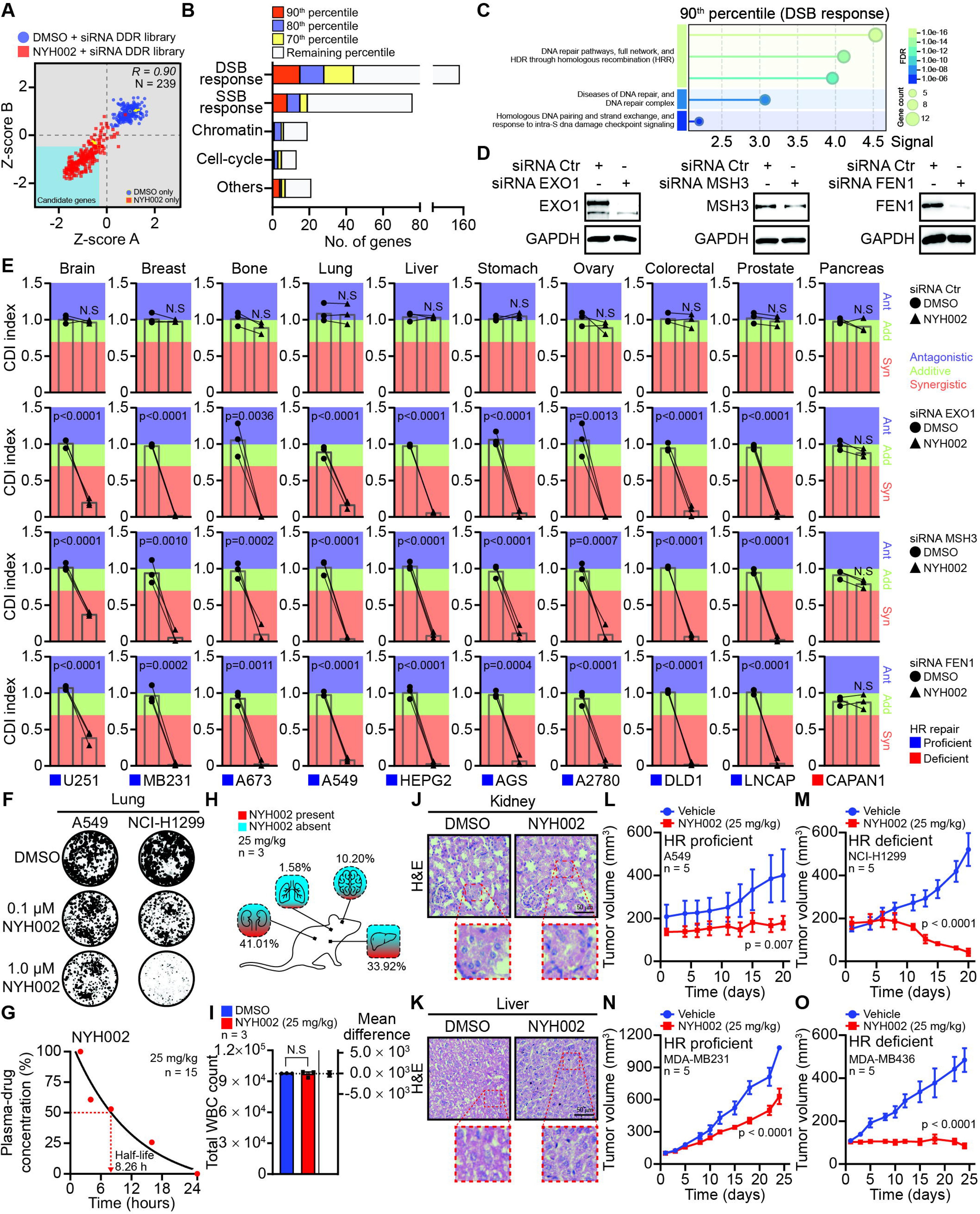
NYH002 therapy is lethal in HR deficient tumors. (A, B) Reproducibility of the siRNA DDR library screen for gene candidates (blue region). Spearman correlation was determined between the two independent replicates of Z-scores A and B in assessing the inhibition of DLD1 cell growth after 72 hours of treatment. The percentile ranking was based on the mean difference in growth inhibition between NYH002- and DMSO-treated DLD1 cells transduced with individual siRNA DDR gene candidates (n = 239). (C) STRING enrichment of the top 90^th^ percentile of gene candidates identified in the DDR response category. (D) siRNA-mediated depletion of HR gene transcripts, identified in the top 90^th^ percentile, was assessed 48 hours post-transfection by immunoblotting WCL of DLD1 cells (E) Synergistic interaction between NYH002 and individual HR genes (*EXO1*, *MSH3*, and *FEN1*) was assessed based on cell survival readout at 72 hours post-treatment and calculated using CDI. Genomic scar scores were selected for nine HR-proficient and one HRD cancer cell line, as indicated. CDIJ<J1 indicates synergistic (Syn), CDIJ=J1 indicates additive (Add), and CDIJ>J1 indicates antagonistic (Ant). (F) Dose response of NYH002 in the colony formation assay of HR-proficient (A549) and HRD (NCI-H1299) cell lines after seven days of treatment (n = 3). (G, H) Pharmacokinetics was determined by intraperitoneal administration of 25 mg/kg NYH002 in BALB/cJ mice, followed by a series of control cull at 2, 4, 8, 16, and 24 hours. Plasma and animal organs were processed and subsequently resuspended in ACN solution to precipitate NYH002 for detection and quantification by MS. (I, J, K) Cytotoxicity of NYH002 was assessed by administering either DMSO or 25 mg/kg NYH002 via intraperitoneal injection into nude mice on alternate days for two weeks, followed by a control cull to collect whole blood for hematologic viability analysis. The kidney and liver were also fixed and processed for H&E staining to assess changes in tissue architecture and the presence of necrotic cells (L, M) Assessment of NYH002 therapeutic efficacy in xenograft lung cancer models. Nude mice were randomly assigned and implanted with 1 x 10^6^ HR-proficient (A549) or HRD (NCI-H1299) cells into the subcutaneous space of the animal’s flank. Treatment was subsequently administered as either vehicle or 25 mg/kg NYH002 via intratumoral injection. Tumor size was measured alternate days for a period of two weeks. A control cull was performed upon completion of the treatment regimen. (N, O) Therapeutic assessment of NYH002 in orthotopic breast cancer models. Nude mice were randomly assigned and implanted with 1 x 10^6^ of HR-proficient (MDA-MB231) and HRD (MDA-MB436) cells in the mammary fat pad. Treatment was subsequently administered as either vehicle or 25 mg/kg NYH002 via intraperitoneal injection. Tumor size was measured every alternate day for two weeks. A control cull was performed upon completion of the treatment regimen. Data are presented as mean ± SEM. Unpaired two-tailed t-test (E and I) and two-way ANOVA (L, M, N and O). p-values < 0.05 were considered significant whereas N.S was regarded as not significant. See also Figure S6

In our preclinical assessment, we identified and selected the most frequent occurring HRD tumor type in the CCLE dataset as a representative model to investigate the therapeutic potential of NYH002 in targeting solid tumors with deficient HR (Figure S6O–P). Based on this analysis, we prioritized on lung and breast cancers. Clonogenic and cell viability assays confirmed that H1299 (lung) and MDA-MB-436 (breast) cells with HRD exhibited significant vulnerability to NYH002 therapy, whereas their HR-proficient counterparts, A549 and MDA-MB-231, exhibited better tolerance under identical treatment conditions (Figure S6Q and 6F). Our dose escalation assay in non-tumor bearing animals determined that 25 mg/kg of NYH002 represents the relative minimum effective dose with complete drug clearance observed by 24 hours (Figure S6R and 6G). Intraperitoneal administration of NYH002 at this concentration led to widespread distribution across all major organs within six hours, as confirmed by LC-MS/MS. Given the molecular weight of NYH002 is approximately 450 Da, the ability of this small molecule to penetrate the blood-brain barrier is consistent with its physicochemical properties (Figure 6H) (40). Importantly, there was no adverse and chronic systemic cytotoxicity even after two weeks of intraperitoneal administration and this finding was corroborated by stable blood counts (Figure 6I) and an absence of cell death, as confirmed by H&E staining of the liver and kidney (Figure 6J–K). Across the four xenograft models encompassing both HRD and HR-proficient subtypes of lung and breast cancer, administration of NYH002 at a dosage of 25 mg/kg consistently reduced tumor growth in all animals (Figure 6L–O). Its therapeutic efficacy, however, was remarkable in HRD tumors, where growth was abrogated, highlighting NYH002’s selective cytotoxicity against malignancies with compromised HR (Figure S6S–T).

## DISCUSSION

R-loop represents a fundamental paradox in tumor malignancy, indispensable for cellular functions, yet lethal in excessive amount (41). Their threat to genome integrity arises when R-loops drive TRCs, a context that motivated the discovery of ILF2’s critical role in safeguarding replication stress-induced DNA DSBs. While replication and transcription are mechanistically distinct and temporally separated processes, they are frequently converged on the same genomic locus. Precise coordination of R-loop resolution is essential to prevent deleterious head-on collisions between transcription machinery and advancing replication forks. The fidelity of DNA replication during active transcription is dependent on ILF2 interaction with various helicases. By direct binding with ILF2, helicases such as DHX9 can actively unwind stable RNA:DNA hybrids (42) and assist RNAPII in engaging template DNA to initiate nascent transcripts synthesis (43, 44). This finding is consistent with our observation in the reduced activities at TSSs and the stalling of RNAPII from unresolved R-loops (45). Like DHX9, other helicases including SETX and WRN also play an immediate role in the unwinding of RNA:DNA hybrid structures (28, 46). BRCA1 and USP11 are known recruiters for SETX (29), analogous to CK2 in activating WRN to initiate R-loop resolution (47). Current evidence is clear that each helicase is subject to distinct regulatory mechanisms, underscoring their independent roles in R-loop resolution. Conversely, emerging data also demonstrated a multitude of these helicases are conserved and can act in concert through direct interaction with each other to ensure timely resolution of R-loop (27). It is possible independent pathways that regulate different helicases are necessary for redundancies, but the understanding of how these different mechanisms cooperate to maintain R-loop homeostasis remain incomplete.

Our work offers an alternative model in which ILF2 overrides these individual pathways by acting as a central regulator. Depletion of ILF2 triggers a cascade of protein destabilization via DHX9, including SETX and WRN, while also impacting members of the DEAD/DEAH-box RNA helicases; DDX3X, DDX5, and DHX15 (48–50). ILF2’s ubiquitous expression and near absence of somatic mutations argue against a classical oncogenic role (51). Instead, ILF2 acts as a non-oncogenic factor, leveraging its upstream regulatory function in R-loop resolution to mitigate genomic stress driven by TRCs.

The exploitation of ILF2’s role as a central regulator of RNA:DNA helicases led to the discovery of NYH002 as a first-in-class ILF2 inhibitor (Figure 7A). NYH002-treated cancer cells phenocopy ILF2 depletion by disrupting the signaling network essential for R-loop resolution. Several helicase inhibitors have been developed (11, 52). Their therapeutic application is however hindered by their essential roles in RNA and DNA metabolism, complicating efforts to selectively target R-loop resolution without disrupting other critical cellular processes. Inhibiting ILF2 as the central regulator coordinates the removal of all downstream RNA:DNA helicase activities dedicated to R-loop resolution and enables a precise targeting without compromising other helicase-associated pathways. Additionally, our PK and biodistribution assessment demonstrated that the serum concentration required for NYH002 to impede tumor progression is readily achievable without chronic adverse side effects. An independent phenotypic study corroborated our findings, confirming that Molephantin exerts selective cytotoxicity against high-grade cancers (53). These results align with the retrospective observational data, where cancer patients undergoing integrative medicine did not exhibit long-term systemic therapeutic toxicity. Instead, routine medical assessments and patient-reported outcomes concluded stable disease or tumor reduction.

**Figure 7:**
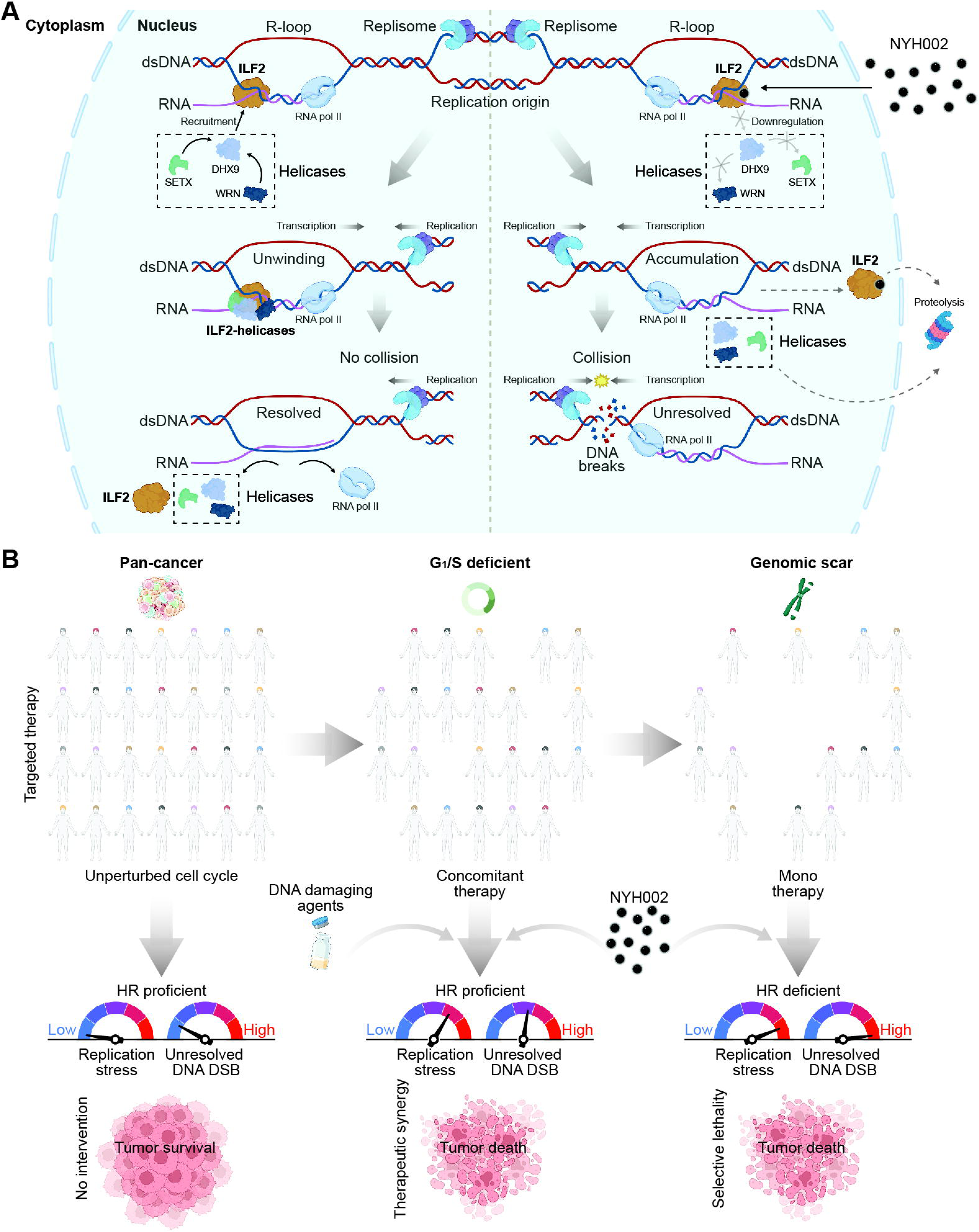
A proposed model of NYH002. (A, B) Therapeutic framework illustrating the mechanisms of action of NYH002. Created with BioRender.com.

The stark contrast in NYH002’s therapeutic cytotoxicity in eliminating tumorigenic cells and leaving minimal harm on healthy tissue can be explained by modeling cell cycle checkpoint progression (Figure 7B). Tumors with G_1_/S deficiency have considerable replication stress (32). When NYH002 disrupts DNA replication by introducing unresolved R-loops, these tumors encounter an increased frequency of TRCs, leading to replication stress-induced DNA breakages. Repair of these S-phase DSBs is initiated by DNA end resection (54), a process that biases toward HR and compels NYH002-treated tumors to rely on this post-replicative DSB repair pathway for survival. We exploited the dependency of HR-mediated repair by combining NYH002 with different DNA damage inducing agents to exacerbate DSB accumulation beyond the threshold of repairability. To further examine this therapeutic strategy, we extended the investigation to uncover vulnerabilities in HRD cancer cells that are independent of BRCA1/2 status to broaden the treatment potential of NYH002 (39). Our data present multiple lines of evidence indicating that the disruption of R-loop homeostasis constitutes a key determinant for selective killing in cancer cells. Here, G_1_/S-deficient cancer cells are more vulnerable to NYH002 in triggering TRCs. Overexpression of RNaseH1 counteracts NYH002-induced R-loop accumulation via RNA strand degradation, thereby rescuing replication stress–induced DNA breakages. The relationship of synthetic lethal HR gene pairs *EXO1*, *FEN1* and *MSH3* in ILF2-depleted cells is substantiated across multiple solid tumor types. Lastly, NYH002 as a monotherapy suppresses tumor growth in both BRCA1/2-independent HR-deficient xenografts and BRCA1/2-mutant orthotopic models. Our findings inform a mechanistic framework that has been mirrored by emerging reports describing two individual synthetic lethality strategies; disrupting transcription factors to elevate R-loop accumulation in DNA repair deficient cells (22) and promoting the displacement of exoribonucleases in cancers burdened by excessive R-loops (55). Collectively, our study positions NYH002 as an actionable therapeutic agent that leverages ILF2’s central role in R-loop resolution to eliminate HRD cancer cells, providing a distinct strategy beyond conventional DNA repair–targeted approaches.

## MATERIAL AND METHODS

### Human and Animal Ethics

Patient recruitments were carried out in accordance with the approval of the Institutional Review Board of Nanyang Technological University (NTU), Singapore (IRB-2022-768). For animal studies, experiments were conducted in accordance with the recommendations of the National Advisory Committee for Laboratory Animal Research guidelines, under the Animal & Birds (Care and Use of Animals for Scientific Purposes) Rules of Singapore. The Institutional Animal Care and Use Committee of NTU approved the current protocol (A22049).

### Observational Study

In a retrospective research initiative conducted offsite at NTU, patients who had previously undergone integrative medicine treatment were invited to participate in an observational study. Participant eligibility was assessed in accordance with the approved protocol (IRB-2022-768). Evaluation was based on a review of participants’ past medical histories, with a specific inclusion criterion stipulating a minimum six months of integrative medicine use. The integrative medicine regimen comprised a minimum daily oral administration of combined PAs, which were derived from medicinal plant materials. Additional criteria included the periodic conduct of questionnaires, and the collection of routine pathological reports of biomarkers and/or medical imaging conducted before and after the integrative medicine treatment. Informed consent was obtained from all patients. All biographical data were de-identified. Financial compensation was not offered for participation in this study.

### Cell Culture

Unless otherwise stated, all cell lines were grown in culture medium supplemented with 10% fetal bovine serum (FBS) (Thermo Fisher Scientific) and 1% penicillin/streptomycin (Cytiva). DLD-1, HCT116, H1299, Huh7, AGS, PC3, A2780, and A673, A549, HepG2, LnCAP cells were cultured in RPMI1640 Medium (Gibco). MDA-MB-231, MDA-MB-436, U2OS, and U251 were cultured in DMEM (Gibco). Capan-1 was cultured in IMDM (Gibco) supplemented with 20% FBS and 1% penicillin/streptomycin. BJ-5Ta was cultured in a 1:1 ratio of DMEM and Medium 199, supplemented with 10% FBS, 1% penicillin/streptomycin, and 0.01 mg/ml hygromycin B (Nacalai Tesque). All cells were cultured in a humidified incubator at 37°C with 5% CO₂, routinely screened for mycoplasma contamination, and authenticated by STR profile.

### Phytotherapeutics

Medicinal plant materials, PA1 (*Clinacanthus nutans*), PA2 (*Strobilanthes crispa L*), PA3 (*Elephantopus tomentosus L*), PA4 (*Leea indica*), PA5 (*Houttuynia cordata*), and PA6 (*Zebrina pendula Schnizl*) were cleaned, cryodesiccated and grounded into a fine powder with a BioloMix grinder. To achieve bioactive extracts, the powdered materials were suspended in 60% aqueous ethanol at a ratio of 1:30 (w/v) and subjected to continuous agitation at room temperature. Mixture was centrifuged, and the resulting supernatant was concentrated overnight at 37°C using a vacuum centrifuge (Concentrator Plus, Eppendorf, USA). The dry pellet was weighed and stored at 4°C. As a ready-to-use bioactive extract, the pellet was resuspended in sterile water.

### Chemicals

APH, RO-3306, MMS and 5FU were purchased from Sigma-Aldrich. TMZ and 5,6-Dichloro-1-β-D-ribofuranosylbenzimidazole (DRB) were obtained from MedChem Express and Merck, respectively.

### RNA Interference

The human ON-TARGET*plus* DDR siRNA library (Dharmacon) was obtained as SMARTpool reagents, each comprising a mixture of four siRNAs targeting a single gene. In total, the library targeted 239 genes (56). Additional ON-TARGET*plus* siRNA was also purchased for *EXO1*, *MSH3*, *FEN1* genes. siRNA negative control was obtained from Dharmacon.

### Plasmid

RNaseH1 plasmid was a kind gift from Dr. Kah Suan Lim, National Cancer Centre, Singapore.

### Public Datasets

Genomic, exon and RNA-seq datasets were obtained from TCGA (57).

### Bioactive Compounds

PA3 crude powder (40 g) was sonicated in ethyl acetate and decanted repeatedly to achieve a greenish-yellow supernatant. The combined crude extracts were then filtered over Celite®, concentrated *in vacuo*, and purified by flash column chromatography (FCC) on silica gel (n-hexane/ethyl acetate, 4:1 to 1:1). Purified material in ethyl acetate was treated with charcoal before passing through Celite®. The filtrate was concentrated under reduced pressure and further purified by gel permeation chromatography (LaboACE LC-5060, JAIGEL-2HR-40 column; 24 mg/ml; 10 ml injection; 30 ml/min flow rate), affording three fractions: Molephantinin and Molephantin, and a mixture of both compounds.

### Molephantin Derivatives and Probe

NYH001 derivatives were synthesized for SAR. Substituted benzoyl chlorides with various functional groups were added to a mixture of NYH001, Et_3_N, and DMAP in anhydrous CH_2_Cl_2_ at 0°C under argon. The reaction was performed at room temperature and concentrated *in vacuo*. The crude residue was further rinsed with saturated NaHCO_3_, extracted with dichloromethane, and the combined organic layers were filtered, purified, and further concentrated using FCC on silica gel to produce solids of various derivatives. The structures and qualities of all compounds were examined using ^1^H NMR, ^13^C NMR, and MS (ESI). Please refer to supplementary data for detailed protocol and characterization reports.

### Cell Divisions

Briefly, 2 × 10^6^ viable single cells were pre-labeled with 10 μM CFSE (Thermo Fisher Scientific). Labeling was quenched with RPMI medium containing 10% FBS. A portion of these cells was fixed and stored at 4°C, while the remaining cells (5 × 10^5^) were seeded in a T-25 flask with the appropriate treatment. After four days of culture, cells were harvested and rinsed twice with PBS prior to flow cytometric analysis. Cell proliferation data were analyzed using FlowJo software.

### Cell Proliferation and Therapeutic Synergy

Cell viability was assessed using MTT. Briefly, cells were seeded in 96-well plates at a density of 8 × 10^3^ to 1.2 × 10^4^ cells per well the day prior to experimentation. Following the indicated treatment, MTT was added to cells and incubated in the dark at 37°C. Formazan crystals were solubilized with DMSO. Absorbance was measured at 570 nm. Cell viability was calculated as a percentage relative to the untreated control. EC_50_ values were calculated using GraphPad. Alternatively, cell viability data were organized into a matrix format for analysis using SynergyFinder 3.0 (58). The Highest Single Agent (HSA) model was selected for its conservative nature, minimizing false positives and reducing the risk of overestimating synergy between two drugs. For synthetic lethality, the coefficient of drug interaction (CDI) was calculated by dividing the observed cell viability of the combined treatment by the product of the cell viabilities for each individual treatment at the same concentration (59).

### Colony Formation

Approximately 1 × 10^3^ cells were seeded into 6-well plates. The cells were treated with 0.1 μM and 1 μM NYH002 for one hour at 37°C and incubated in fresh media for seven days to allow colony formation. The colonies were fixed in 4% paraformaldehyde (PFA) and stained with 0.5% crystal violet. Colonies were rinsed with water and air-dried, Images were taken with ChemiDoc MP Imaging System (Bio-Rad). The number and area of colonies were quantified using FIJI software.

### Cell Cycle Profile

Approximately 6 × 10^5^ cells were seeded into 6-well plates and treated with NYH002 for 24 hours. Cells were harvested, fixed in -20°C 70% ethanol and stained with propidium iodide staining buffer (0.1% Triton-X, 40 mg/ml propidium iodide, 200 mg/ml RNase A). Samples were quantified with LSRFortessa™ (BD Biosciences) and analyzed in FlowJo™ software.

### Immunofluorescence Stain

Cells were seeded onto coverslips. Following treatment the next day, cells were fixed with 4% PFA and permeabilized with 0.2% Triton X-100/ PBS. For S9.6 staining, cells were fixed with ice-cold methanol. Primary antibodies were applied at room temperature, rinsed with TBS/0.1% Tween, and replaced with Alexa Fluor-conjugated secondary antibodies. Nuclei were counterstained with DAPI (1 μg/ml), and coverslips were mounted in ProLong™ Gold (Thermo Fisher Scientific). The slides were imaged using a Zeiss Axio Observer.Z1 fluorescence microscope or LSM980 confocal microscope. Foci counts were analyzed per nucleus, with at least 100 nuclei per condition analyzed by CellProfiler. Manual verification was performed to exclude artifacts.

### ssDNA Stain

Naïve ssDNA labeling was performed as previously described (60). Briefly, cells were pulse-labeled with 10 μM IdU for 24 hours prior to treatment, washed with PBS, and pre-extracted with ice-cold CSK buffer (10 mM PIPES, 300 mM sucrose, 100 mM NaCl, 3 mM MgCl_2_, 0.2% Triton X-100) at room temperature. Cells were then fixed in 2% PFA, blocked in 10% BSA in PBS/0.05% Tween and immunostained with anti-IdU primary antibody and Alexa Fluor 594–conjugated secondary antibody.

### DNA Pulse-Labeled Fiber

DNA fiber assay was performed as previously described (61). Briefly, cells were pulse-labeled with 20 μM CIdu (Sigma-Aldrich) for 30 minutes at 37°C, treated with NYH002 for one hour before labeling with 100 μM IdU (Sigma-Aldrich) for 30 minutes. Cells were then lysed with spreading buffer (0.5% SDS, 200 mM Tris-HCl, 50 mM EDTA) and DNA fibers were stretched on the glass slide. Incorporated analogs were stained using rat anti-BrdU antibody (Abcam) and mouse anti-BrdU antibody (BD Biosciences). Images were acquired with a Zeiss Axio Observer.Z1 fluorescence microscope and at least 200 fibers were quantified for each experiment.

### Chromosome Spread

Treated cells were harvested and treated in hypotonic buffer (75 mM KCl) at 37°C. The buffer was removed by centrifugation and cells were fixed dropwise with ice-cold methanol/acetic acid (3:1). The cells were spun down at 200× g with the fixation step repeated twice before storing samples at 4°C overnight. The cells were dropped onto glass slides and stained with DAPI (1 μg/ml). Images of the chromosome spread were acquired with Zeiss Axio ObserverZ1 fluorescence microscope.

### Cell Transfection

Transfection of RNaseH1 plasmid was performed using Lipofectamine 3000 (Thermo Fisher Scientific) while siRNAs were transfected into cells with DharmaFECT1 (Dharmacon).

### Proximity Ligation Assay

After the indicated treatments, cells cultured on coverslips were fixed with 4% formaldehyde and permeabilized with 0.2% Triton X-100/PBS. PLA was performed using Naveni^TM^ TriFlex Cell MR (Navinvi Diagnostics) according to the manufacturer’s instructions. Briefly, cells were immersed in blocking buffer at 37 °C, then incubated overnight at 4 °C with primary antibodies (1:500) diluted in the supplied buffer. Cells were rinsed and incubated with secondary antibodies conjugated with either plus or minus PLA probes. This is followed by ligation and amplification steps. Cells were stained with DAPI before mounting on glass slides. PLA signal was detected using the Zeiss LSM980 confocal microscope under 40× objective lens. The images were analyzed and processed using Zen Blue software. The PLA signal and foci were quantified using the segmentation tool. Automated segmentation parameters were optimized to detect individual PLA foci. The total number of foci per nucleus was calculated, and at least 100 nuclei per condition was analyzed.

### Cellular Thermal Shift Assay

CETSA was performed following the published protocol (24). Briefly, 1 × 10^7^ DLD-1 cells were resuspended in PBS supplemented with Halt™ Protease and Phosphatase Inhibitor Cocktail (Thermo Fisher Scientific). Cells were subjected to freeze-thaw cycles, and lysate was centrifuged at 12,000× g. The supernatant was collected, aliquoted, and incubated with either DMSO or 100 μM NYH002 at room temperature on a rotator. Samples were subjected to a temperature gradient (40–60°C) by using a gradient thermal cycler (Eppendorf) and subsequently incubated at 25°C before placing on ice. Lysates were centrifuged at 16,000× g, and the supernatant was collected. Prior to SDS-PAGE, Laemmli buffer was added, and samples were denatured at 95°C.

### Cell Lysis and Immunoprecipitation

Cell pellets were lysed in ice-cold RIPA or co-immunoprecipitation lysis buffer and centrifuged at 14,000× g to collect supernatants for immunoblotting and immunoprecipitation, respectively. For the latter, WCL were pre-cleared with Dynabeads™ M-280 streptavidin or protein A (Invitrogen) prior to protein estimation with Pierce BCA Kit (Thermo Fisher Scientific). Briefly, 10 μM of NYH002P was incubated with 1.5 mg of WCL overnight at 4°C, followed by click chemistry-mediated biotinylation and subsequent incubation with Dynabeads™ streptavidin. Alternatively, 1 μM of antibody was incubated with 1 mg of WCL overnight at 4°C and incubated with Dynabeads™ protein A. Samples were magnetically bounded and washed with lysis buffer and a high-salt (0.5 M NaCl) solution. Bound proteins were eluted with 2× Laemmli buffer (Bio-Rad) at 95°C.

### SDS-PAGE, Silver Stain and Immunoblot

Denatured proteins were loaded onto an SDS-PAGE gel and electrophoresed. Silver staining was performed according to the manufacturer’s protocol (Pierce™ Silver Stain Kit). Alternatively, proteins were then transferred onto methanol-activated PVDF membranes using Trans-Blot® Turbo™ System (Bio-Rad). Membranes were blocked with 5% skimmed milk in 0.1% TBS/T prior to incubation with primary antibodies overnight at 4 °C. The following day, membranes were rinsed with 0.1% TBS/T, followed by incubation with horseradish peroxidase (HRP)-conjugated secondary antibodies. Signal detection was performed using enhanced chemiluminescence reagents and visualized with the ChemiDoc MP Imaging System (Bio-Rad).

### Compound Probe-ClickiT

NYH002P was labeled and stained using the Click-iT™ EdU Cell Proliferation Kit for Imaging, Alexa Fluor™ 488 dye (Thermo Fisher Scientific) following manufacturer’s instructions. DLD-1 were seeded and treated with 1 μM NYH002P for six hours. Cells were fixed with 4% PFA and permeabilized with 0.2% Triton-X. Each coverslip was stained with freshly prepared buffer as instructed by the kit and were compatible for further staining with primary antibodies, DAPI, or for PLA experiments.

### RT-qPCR

Total RNA was extracted using FastPure Cell/Tissue Total RNA Isolation Kit V2 (Cat# RC112-01, Vazyme). Total RNA was reverse transcribed to cDNA using HiScript III All-in-one RT SuperMix Perfect for qPCR (Vazyme) on a gradient thermal cycler (Eppendorf). qPCR was performed with 500 nM forward and reverse primers (IDT) and Taq Pro Universal SYBR qPCR Master Mix (Vazyme) using Bio-Rad CFX96 Touch Real-Time PCR Detection System with CFX Manager software (Bio-Rad). *HPRT* was used as an internal control for normalization, and the relative expression level was calculated by the 2(-Delta Delta C(T)) method. Primer sequences are listed in the supplementary data.

### RNA Sequencing

The mRNA of treated samples was purified using poly-T oligo-attached magnetic beads and fragmented for cDNA synthesis. Directional and non-directional libraries were prepared with end repair, A-tailing, adapter ligation, size selection, amplification, and purification. Libraries were quantified and sequenced on an Illumina platform. Quality control included adapter trimming, removal of low-quality reads, and calculation of Q20, Q30, and GC content. Reads were mapped to the reference genome using Hisat2, and novel transcripts were assembled with StringTie. Gene expression levels were quantified using FeatureCounts and FPKM. Differential expression analysis was conducted with DESeq2 using the Benjamini-Hochberg method for multiple testing correction. Genes with an adjusted p-value ≤ 0.05 and absolute fold-change ≥ 2 were considered significantly differentially expressed.

### Proteomics Sample Preparation and Analysis

For each condition, 100 μg of lysate was reduced, alkylated, and incubated overnight with Pierce trypsin protease MS grade (Thermo Fisher Scientific). Label-free or TMT10-plex isobaric-tagged (Thermo Fisher Scientific) samples were first loaded onto an Xbridge™ C18 column where peptides were separated based on their hydrophobicity. The separated peptides were delivered through the Dionex EASY-Spray nano-electrospray to be ionized. These ions were introduced into a high-resolution Q Exactive orbitrap. A full MS scan (350–1,600Jm/z range) was acquired at a resolution of 70,000 and a maximum ion accumulation time of 100 ms. Raw data files were processed and searched using Proteome Discoverer 2.1 (Thermo Fisher Scientific). SEQUEST was then used for protein identification. The normalization mode was based on the total peptide amount and a false discovery rate (FDR) for protein identification was < 1%. The protein list was further filtered with a minimum of 1 unique and 1 common peptide including matched molecular weight prior to pathway enrichment using KEGG, GO, Reactome and WikiPathway by Cytoscape (62). Alternatively, curated proteins were also performed using STRING and IntAct for PPI or the use of CORUM database to match protein complex (23).

### Orthotopic and Xenograft Models

Briefly, tumor cells (1 × 10^6^) in HBSS were injected subcutaneously into the flank or orthotopically into the mammary fat pad of nude mice. Tumors were allowed to grow till they reached a palpable size of approximately 100 mm^3^ before randomizing into control and treated groups. NYH001, NYH002 and NYH003 were dissolved in a vehicle solution consisting of 10% Kolliphor® EL, 10% DMSO, and 80% saline, therapeutics were administered via either the intraperitoneal or intratumoral route. Mice received treatment every other day for two weeks, unless stated otherwise. TMZ was dissolved in saline to a concentration of 10 mg/kg and administered orally once daily. Mice were weighed twice weekly, and tumor dimensions were regularly measured using calipers. Tumor masses were calculated as follows: Tumor volume (mm^3^) = (length × width^2^) / 2. Mice were euthanized in a carbon dioxide chamber when tumor volume reached 1,800 mm^3^, weight loss exceeded 20%, or the treatment duration reached two weeks, whichever occurred first.

### Cytotoxicity, Pharmacokinetics, and Biodistribution

For cytotoxicity assessment, J:NU/Inv mice received intraperitoneal injection of vehicle or 25 mg/kg NYH002 on alternate days for two weeks. After which, blood was collected by cardiac puncture and diluted in PBS containing 1% FBS and 0.1 mM EDTA at a 1:20 ratio, followed by filtration through a 0.45 μm cell strainer and collected in FACS tubes for flow cytometry assessment by cell size and granularity. For pharmacokinetics and biodistribution, BALB/cJ mice received a single intraperitoneal injection of either vehicle or 25 mg/kg NYH002. Blood was collected via cardiac puncture at indicated time points (0 – 24 hours) and transferred into vacutainer^®^ EDTA tubes (BD). Organs were also harvested and homogenized in parallel. Both sample types were subsequently centrifuged. Serum from blood or supernatant from homogenized tissue was collected and individually mixed with ice-cold acetonitrile (ACN) at a 1:3 ratio. Samples were precipitated at −20 °C overnight. The following day, samples were centrifuged at 4°C, and the supernatant was collected and concentrated. Resulting pellet was resuspended in 50% (w/v) ACN and analyzed by MS.

### Histopathological staining

Tissues were fixed in 4% PFA, processed and embedded in paraffin, and subsequently sectioned at 5 μm thickness using a rotary microtome. Deparaffinized samples were stained with Mayer’s hematoxylin and Eosin Y, then dehydrated with ethanol. The slides were mounted with DPX Mountant for histology (Sigma) and imaged using Axio ObserverZ1 fluorescence microscope (Zeiss) under 20× objective lens.

### Hi-C sequencing

Hi-C was performed using the Dovetail TopoLink V2 kit. Libraries were sent to NovogeneAIT for sequencing on Illumina NovaSeq at 300M PE150 reads per sample, with 2 biological replicates per condition. Fastq files from Illumina sequencing were first trimmed with trimmomatic [settings: ILLUMINACLIP: Adapters:2:30:10:8:true \ HEADCROP:5 \ SLIDINGWINDOW:4:10 \ LEADING:3 \ TRAILING:3 \ MINLEN:30]. Reads were aligned on the hg19 genome using bwa default settings. Pairtools was used to prepare the paired interaction using the parse [settings: --min-mapq 10 --walks-policy 5unique –max-inter-align-gap 30], sort [settings: default], dedup [settings: --mark-dups] and split [settings: default] followed by samtools indexing and sorting of resulting .bam files and pairix indexing of resulting .pairs files. Finally, contact matrices were generated using cooler followed by cload [settings: pairix] and zoomify [settings: --balance] commands (63). Pile-ups were generated from .cool contact matrices using coolpup.py and its associated plotpup.py packages (64). The merge matrices from both replicates of each condition were used for this analysis, as it is not influenced by coverage biases. Hi-C heatmaps were generated from the .cool matrices using HiCExplorer (65). The two replicates with the closest coverage quality (U2 and H2 from the shared matrices), at 32 kb resolution, were used for imaging to alleviate coverage biases.

### Illustration

Proposed model for lead compound NYH002 was created with BioRender.com under publication license (MK289PVV3V).

### Data Analysis

All data presented were obtained from three independent experiments, unless otherwise specified. Significance was assessed using an unpaired, two-tailed Student’s t-test, assuming equal variance between the control and experimental groups, when the sample size was 10 or fewer. In the event of a larger sample size (>10), a normality distribution test was conducted using the Shapiro-Wilk Test and QQ-plot. For normally distributed data, a parametric test was used (unpaired two-tailed Student’s t-test or two-way ANOVA test). If the data is not normally distributed, the Mann-Whitney U test was employed. The significance of survival probability was calculated using the log-rank (Mantel-Cox) test. A p-value equal to or less than 0.05 was considered statistically significant in supporting the alternative hypothesis, unless stated otherwise.

## RESOURCE AVAILABILITY

### Lead Contact

Further information and requests for resources should be directed to and will be fulfilled by the lead contact, A/Prof. Li Hoi Yeung.

### Materials Availability

All resources are available upon request to the lead contact.

## ACKNOWLEDGEMENTS

We would like to thank A/Prof. Mu Yuguang and Asst Prof. Wilson Goh for their assistance in molecular docking and MS, respectively; Dr Ming Shan and Chermaine Choo for facilitating patient interviews. This study was supported by the Ministry of Education, Singapore (AcRF Tier 1 RG33/23 to H.Y.L), Nanyang Biologics Pte Ltd (REQ0196946 to H.Y.L and C.G.K) and National Research Foundation, Singapore (MOH-001387 and AISG3-GV-2023-014 to M.J.F)

## AUTHOR CONTRIBUTIONS

Conceptualization, Y.C.L., S.K.L., C.G.K., and H.Y.L.; Formal analysis, Y.C.L., S.K.L., R.H.Y., Y.K.C., B.L., and H.Y.L.; Investigation and data analysis, Y.C.L., S.K.L., R.H.Y., V.M., C.X.C.S., Y.K.C., B.L., M.W., X.M.D., S.K.T., S.L.Y., R.P., S.Z., J.Y.X.W., and Z.A; Resources, P.S., C.N., M.M., S.C., and M.J.F., and L.L.D.Z.; Supervision, Y.C.L., S.K.L., C.G.K., and H.Y.L.; Visualization, Y.C.L., S.K.L., and R.H.Y.; Writing of manuscript, Y.C.L., S.K.L., R.H.Y., and H.Y.L.; Commented on manuscript, all authors; Funding acquisition, M.J.F., C.G.K., and H.Y.L.

## REFERENCE

1. Lai WKM, Pugh BF. Understanding nucleosome dynamics and their links to gene expression and DNA replication. Nat Rev Mol Cell Biol. 2017;18(9):548–62.

2. Kotsantis P, Silva LM, Irmscher S, Jones RM, Folkes L, Gromak N, et al. Increased global transcription activity as a mechanism of replication stress in cancer. Nat Commun. 2016;7:13087.

3. Roy D, Zhang Z, Lu Z, Hsieh CL, Lieber MR. Competition between the RNA transcript and the nontemplate DNA strand during R-loop formation in vitro: a nick can serve as a strong R-loop initiation site. Mol Cell Biol. 2010;30(1):146–59.

4. Hamperl S, Bocek MJ, Saldivar JC, Swigut T, Cimprich KA. Transcription-Replication Conflict Orientation Modulates R-Loop Levels and Activates Distinct DNA Damage Responses. Cell. 2017;170(4):774–86 e19.

5. Lam FC, Kong YW, Huang Q, Vu Han TL, Maffa AD, Kasper EM, et al. BRD4 prevents the accumulation of R-loops and protects against transcription-replication collision events and DNA damage. Nat Commun. 2020;11(1):4083.

6. Bayona-Feliu A, Herrera-Moyano E, Badra-Fajardo N, Galvan-Femenia I, Soler-Oliva ME, Aguilera A. The chromatin network helps prevent cancer-associated mutagenesis at transcription-replication conflicts. Nat Commun. 2023;14(1):6890.

7. Patel PS, Algouneh A, Krishnan R, Reynolds JJ, Nixon KCJ, Hao J, et al. Excessive transcription-replication conflicts are a vulnerability of BRCA1-mutant cancers. Nucleic Acids Res. 2023;51(9):4341–62.

8. Shinriki S, Hirayama M, Nagamachi A, Yokoyama A, Kawamura T, Kanai A, et al. DDX41 coordinates RNA splicing and transcriptional elongation to prevent DNA replication stress in hematopoietic cells. Leukemia. 2022;36(11):2605–20.

9. Gan W, Guan Z, Liu J, Gui T, Shen K, Manley JL, et al. R-loop-mediated genomic instability is caused by impairment of replication fork progression. Genes Dev. 2011;25(19):2041–56.

10. Safari M, Litman T, Robey RW, Aguilera A, Chakraborty AR, Reinhold WC, et al. R-Loop-Mediated ssDNA Breaks Accumulate Following Short-Term Exposure to the HDAC Inhibitor Romidepsin. Mol Cancer Res. 2021;19(8):1361–74.

11. Manzo SG, Hartono SR, Sanz LA, Marinello J, De Biasi S, Cossarizza A, et al. DNA Topoisomerase I differentially modulates R-loops across the human genome. Genome Biol. 2018;19(1):100.

12. Piekarz RL, Frye R, Prince HM, Kirschbaum MH, Zain J, Allen SL, et al. Phase 2 trial of romidepsin in patients with peripheral T-cell lymphoma. Blood. 2011;117(22):5827–34.

13. Wagener DJ, Verdonk HE, Dirix LY, Catimel G, Siegenthaler P, Buitenhuis M, et al. Phase II trial of CPT-11 in patients with advanced pancreatic cancer, an EORTC early clinical trials group study. Ann Oncol. 1995;6(2):129–32.

14. Hosea R, Hillary S, Naqvi S, Wu S, Kasim V. The two sides of chromosomal instability: drivers and brakes in cancer. Signal Transduct Target Ther. 2024;9(1):75.

15. Tutt ANJ, Garber JE, Kaufman B, Viale G, Fumagalli D, Rastogi P, et al. Adjuvant Olaparib for Patients with BRCA1- or BRCA2-Mutated Breast Cancer. N Engl J Med. 2021;384(25):2394–405.

16. Yap TA, O’Carrigan B, Penney MS, Lim JS, Brown JS, Luken MJdM, et al. Phase I Trial of First-in-Class ATR Inhibitor M6620 (VX-970) as Monotherapy or in Combination With Carboplatin in Patients With Advanced Solid Tumors. J Clin Oncol. 2020;38:3195–204.

17. Patouret R, Cham N, Chiba S. Collective Synthesis of Highly Oxygenated (Furano)germacranolides Derived from Elephantopus mollis and Elephantopus tomentosus. Angew Chem Int Ed Engl. 2024;63(19):e202402050.

18. Jung Y, Kraikivski P, Shafiekhani S, Terhune SS, Dash RK. Crosstalk between Plk1, p53, cell cycle, and G2/M DNA damage checkpoint regulation in cancer: computational modeling and analysis. NPJ Syst Biol Appl. 2021;7(1):46.

19. Leriche M, Bonnet C, Jana J, Chhetri G, Mennour S, Martineau S, et al. 53BP1 interacts with the RNA primer from Okazaki fragments to support their processing during unperturbed DNA replication. Cell Rep. 2023;42(11):113412.

20. Lukas C, Savic V, Bekker-Jensen S, Doil C, Neumann B, Pedersen RS, et al. 53BP1 nuclear bodies form around DNA lesions generated by mitotic transmission of chromosomes under replication stress. Nat Cell Biol. 2011;13(3):243–53.

21. Einig E, Jin C, Andrioletti V, Macek B, Popov N. RNAPII-dependent ATM signaling at collisions with replication forks. Nat Commun. 2023;14(1):5147.

22. Awwad SW, Doyle C, Coulthard J, Bader AS, Gueorguieva N, Lam S, et al. KLF5 loss sensitizes cells to ATR inhibition and is synthetic lethal with ARID1A deficiency. Nat Commun. 2025;16(1):480.

23. Tsitsiridis G, Steinkamp R, Giurgiu M, Brauner B, Fobo G, Frishman G, et al. CORUM: the comprehensive resource of mammalian protein complexes-2022. Nucleic Acids Res. 2023;51(D1):D539-D45.

24. Jafari R, Almqvist H, Axelsson H, Ignatushchenko M, Lundback T, Nordlund P, et al. The cellular thermal shift assay for evaluating drug target interactions in cells. Nat Protoc. 2014;9(9):2100–22.

25. Chiu CL, Li CG, Verschueren E, Wen RM, Zhang D, Gordon CA, et al. NUSAP1 Binds ILF2 to Modulate R-Loop Accumulation and DNA Damage in Prostate Cancer. Int J Mol Sci. 2023;24(7).

26. Cristini A, Groh M, Kristiansen MS, Gromak N. RNA/DNA Hybrid Interactome Identifies DXH9 as a Molecular Player in Transcriptional Termination and R-Loop-Associated DNA Damage. Cell Rep. 2018;23(6):1891–905.

27. Petermann E, Lan L, Zou L. Sources, resolution and physiological relevance of R-loops and RNA-DNA hybrids. Nat Rev Mol Cell Biol. 2022;23(8):521–40.

28. Marabitti V, Lillo G, Malacaria E, Palermo V, Sanchez M, Pichierri P, et al. ATM pathway activation limits R-loop-associated genomic instability in Werner syndrome cells. Nucleic Acids Res. 2019;47(7):3485–502.

29. Rao S, Andrs M, Shukla K, Isik E, Konig C, Schneider S, et al. Senataxin RNA/DNA helicase promotes replication restart at co-transcriptional R-loops to prevent MUS81-dependent fork degradation. Nucleic Acids Res. 2024;52(17):10355–69.

30. Kitange GJ, Mladek AC, Schroeder MA, Pokorny JC, Carlson BL, Zhang Y, et al. Retinoblastoma Binding Protein 4 Modulates Temozolomide Sensitivity in Glioblastoma by Regulating DNA Repair Proteins. Cell Rep. 2016;14(11):2587–98.

31. Martino-Echarri E, Henderson BR, Brocardo MG. Targeting the DNA replication checkpoint by pharmacologic inhibition of Chk1 kinase: a strategy to sensitize APC mutant colon cancer cells to 5-fluorouracil chemotherapy. Oncotarget. 2014;5:9889–900.

32. Kok YP, Guerrero Llobet S, Schoonen PM, Everts M, Bhattacharya A, Fehrmann RSN, et al. Overexpression of Cyclin E1 or Cdc25A leads to replication stress, mitotic aberrancies, and increased sensitivity to replication checkpoint inhibitors. Oncogenesis. 2020;9(10):88.

33. Khanna KK, Keating KE, Kozlov S, Scott S, Gatei M, Hobson K, et al. ATM associates with and phosphorylates p53: mapping the region of interaction. Nat Gen. 1998;20:398–400.

34. Guo E, Ishii Y, Mueller J, Srivatsan A, Gahman T, Putnam CD, et al. FEN1 endonuclease as a therapeutic target for human cancers with defects in homologous recombination. Proc Natl Acad Sci U S A. 2020;117(32):19415–24.

35. Li Y, Zhang Y, Shah SB, Chang CY, Wang H, Wu X. MutSbeta protects common fragile sites by facilitating homology-directed repair at DNA double-strand breaks with secondary structures. Nucleic Acids Res. 2024;52(3):1120–35.

36. van de Kooij B, Schreuder A, Pavani R, Garzero V, Uruci S, Wendel TJ, et al. EXO1 protects BRCA1-deficient cells against toxic DNA lesions. Mol Cell. 2024;84(4):659–74 e7.

37. Nguyen L, J WMM, Van Hoeck A, Cuppen E. Pan-cancer landscape of homologous recombination deficiency. Nat Commun. 2020;11(1):5584.

38. Barretina J, Caponigro G, Stransky N, Venkatesan K, Margolin AA, Kim S, et al. The Cancer Cell Line Encyclopedia enables predictive modelling of anticancer drug sensitivity. Nature. 2012;483(7391):603–7.

39. Rempel E, Kluck K, Beck S, Ourailidis I, Kazdal D, Neumann O, et al. Pan-cancer analysis of genomic scar patterns caused by homologous repair deficiency (HRD). NPJ Precis Oncol. 2022;6(1):36.

40. Wu D, Chen Q, Chen X, Han F, Chen Z, Wang Y. The blood-brain barrier: structure, regulation, and drug delivery. Signal Transduct Target Ther. 2023;8(1):217.

41. Garcia-Muse T, Aguilera A. R Loops: From Physiological to Pathological Roles. Cell. 2019;179(3):604–18.

42. Chakraborty P, Grosse F. Human DHX9 helicase preferentially unwinds RNA-containing displacement loops (R-loops) and G-quadruplexes. DNA Repair. 2011;10(6):654–65.

43. Chakraborty P, Huang JTJ, Hiom K. DHX9 helicase promotes R-loop formation in cells with impaired RNA splicing. Nat Commun. 2018;9(1):4346.

44. Yang BZ, Liu MY, Chiu KL, Chien YL, Cheng CA, Chen YL, et al. DHX9 SUMOylation is required for the suppression of R-loop-associated genome instability. Nat Commun. 2024;15(1):6009.

45. Zardoni L, Nardini E, Brambati A, Lucca C, Choudhary R, Loperfido F, et al. Elongating RNA polymerase II and RNA:DNA hybrids hinder fork progression and gene expression at sites of head-on replication-transcription collisions. Nucleic Acids Res. 2021;49(22):12769–84.

46. Skourti-Stathaki K, Proudfoot NJ, Gromak N. Human senataxin resolves RNA/DNA hybrids formed at transcriptional pause sites to promote Xrn2-dependent termination. Mol Cell. 2011;42(6):794–805.

47. Noto A, Valenzisi P, Di Feo F, Fratini F, Kulikowicz T, Sommers JA, et al. Phosphorylation-dependent WRN-RPA interaction promotes recovery of stalled forks at secondary DNA structure. Nat Commun. 2025;16(1):997.

48. Polenkowski M, Allister AB, Burbano de Lara S, Pierce A, Geary B, El Bounkari O, et al. THOC5 complexes with DDX5, DDX17, and CDK12 to regulate R loop structures and transcription elongation rate. iScience. 2023;26(1):105784.

49. Secchi M, Garbelli A, Riva V, Deidda G, Santonicola C, Formica TM, et al. Synergistic action of human RNaseH2 and the RNA helicase-nuclease DDX3X in processing R-loops. Nucleic Acids Res. 2024;52(19):11641–58.

50. Yu Z, Mersaoui SY, Guitton-Sert L, Coulombe Y, Song J, Masson JY, et al. DDX5 resolves R-loops at DNA double-strand breaks to promote DNA repair and avoid chromosomal deletions. NAR Cancer. 2020;2(3):zcaa028.

51. Uhlen M, Fagerberg L, Hallstrom BM, Lindskog C, Oksvold P, Mardinoglu A, et al. Proteomics. Tissue-based map of the human proteome. Science. 2015;347(6220):1260419.

52. Nguyen GH, Dexheimer TS, Rosenthal AS, Chu WK, Singh DK, Mosedale G, et al. A small molecule inhibitor of the BLM helicase modulates chromosome stability in human cells. Chem Biol. 2013;20(1):55–62.

53. Ling Z, Pan J, Zhang Z, Chen G, Geng J, Lin Q, et al. Small-molecule Molephantin induces apoptosis and mitophagy flux blockage through ROS production in glioblastoma. Cancer Lett. 2024;592:216927.

54. Bolderson E, Tomimatsu N, Richard DJ, Boucher D, Kumar R, Pandita TK, et al. Phosphorylation of Exo1 modulates homologous recombination repair of DNA double-strand breaks. Nucleic Acids Res. 2010;38(6):1821–31.

55. Krishnan R, Lapierre M, Gautreau B, Nixon KCJ, El Ghamrasni S, Patel PS, et al. RNF8 ubiquitylation of XRN2 facilitates R-loop resolution and restrains genomic instability in BRCA1 mutant cells. Nucleic Acids Res. 2023;51(19):10484–505.

56. Barroso S, Herrera-Moyano E, Munoz S, Garcia-Rubio M, Gomez-Gonzalez B, Aguilera A. The DNA damage response acts as a safeguard against harmful DNA-RNA hybrids of different origins. EMBO Rep. 2019;20(9):e47250.

57. Ciriello G, Miller ML, Aksoy BA, Senbabaoglu Y, Schultz N, Sander C. Emerging landscape of oncogenic signatures across human cancers. Nat Genet. 2013;45(10):1127–33.

58. Ianevski A, Giri AK, Aittokallio T. SynergyFinder 3.0: an interactive analysis and consensus interpretation of multi-drug synergies across multiple samples. Nucleic Acids Res. 2022;50(W1):W739–W43.

59. Kim H, Xu H, George E, Hallberg D, Kumar S, Jagannathan V, et al. Combining PARP with ATR inhibition overcomes PARP inhibitor and platinum resistance in ovarian cancer models. Nat Commun. 2020;11(1):3726.

60. Lim YC, Ensbey KS, Offenhauser C, D’Souza R CJ, Cullen JK, Stringer BW, et al. Simultaneous targeting of DNA replication and homologous recombination in glioblastoma with a polyether ionophore. Neuro Oncol. 2020;22(2):216–28.

61. Maya-Mendoza A, Moudry P, Merchut-Maya JM, Lee M, Strauss R, Bartek J. High speed of fork progression induces DNA replication stress and genomic instability. Nature. 2018;559(7713):279–84.

62. Bindea G, Mlecnik B, Hackl H, Charoentong P, Tosolini M, Kirilovsky A, et al. ClueGO: a Cytoscape plug-in to decipher functionally grouped gene ontology and pathway annotation networks. Bioinformatics. 2009;25(8):1091–3.

63. Abdennur N, Mirny LA. Cooler: scalable storage for Hi-C data and other genomically labeled arrays. Bioinformatics. 2020;36(1):311–6.

64. Flyamer IM, Illingworth RS, Bickmore WA. Coolpup.py: versatile pile-up analysis of Hi-C data. Bioinformatics. 2020;36(10):2980–5.

65. Wolff J, Rabbani L, Gilsbach R, Richard G, Manke T, Backofen R, et al. Galaxy HiCExplorer 3: a web server for reproducible Hi-C, capture Hi-C and single-cell Hi-C data analysis, quality control and visualization. Nucleic Acids Res. 2020;48(W1):W177–W84.

