## Supplementary figure for "ILF2 blockade suppresses helicase-mediated R-loop resolution to induce lethality in homologous recombination-deficient cancers"

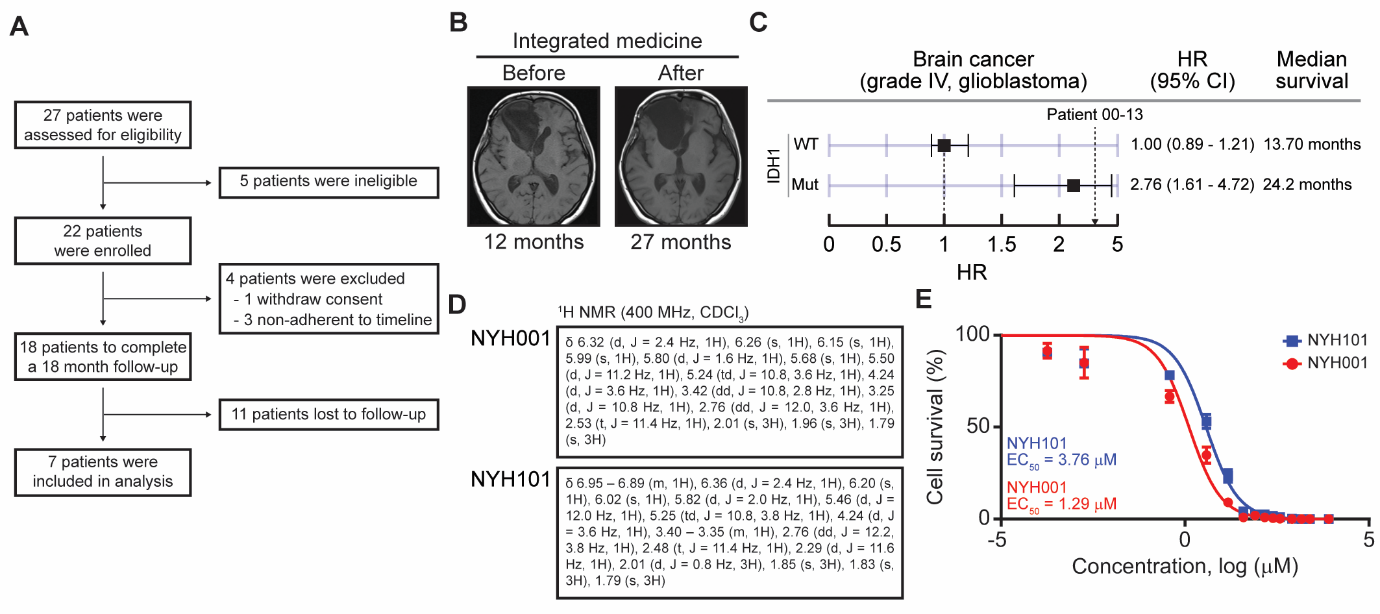

**Supplementary Figure 1:** Discovery of NYH001 and NYH101 compounds in patient cohort receiving integrative medicine.

(A) Flowchart of patient recruitment for eligibility and inclusion in the retrospective observational study.

(B) T1-weighted MRI scans of patient 00-13 before and after receiving integrated therapy.

(C) Hazard ratio (HR) of glioblastoma patients from the TCGA database in comparison to the existing survival timeline (alive) of patient 00-13.

(D) ^1^H NMR structural information of the naturally occurring NYH001 (Molephantin) and NYH101 (Molephantinin).

(E) Dose response of NYH001 and NYH101 in DLD1 cells at 72 hours post-treatment (n = 3).

Data are presented as mean ± SEM.

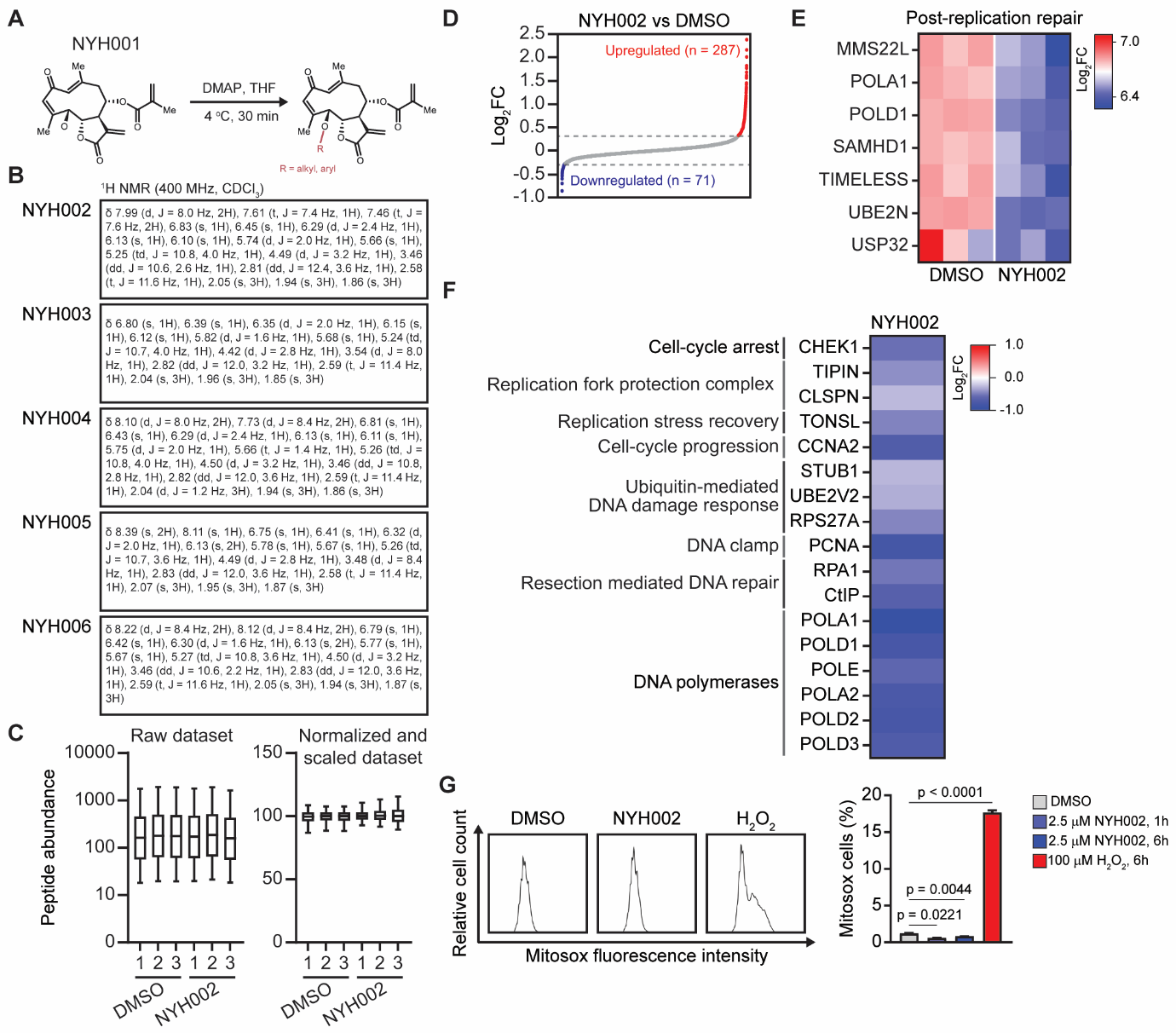

**Supplementary Figure 2:** Lead-optimized NYH001 suppresses the replication machinery leading to chromosomal aberrations.

(A) An overview of the strategy used for the derivatization of Molephantin molecules.

(B) ^1^H NMR structural information of Molephantin derivatives (NYH002 – NYH006).

(C, D) MS of TMT-labeled WCL from DLD1 cells. Raw peptide abundances from independent runs were normalized and scaled. The mean value of each identified protein was then converted to log_2_FC and presented as a waterfall plot.

(E) Differential expression of proteins that are enriched from the “post-replication repair” pathway.

(F) Heatmap of downregulated transcripts used to identify secondary clusters in the STRING PPI network.

(G) Oxidative stress was assessed by staining live DLD1 cells with MitoSOX and analyzing the indicated samples via flow cytometry (n = 3). Treatment with 100 µM H_2_O_2_ for six hours was used as a positive control (n = 3).

Data are presented as mean ± SEM. p-values < 0.05 were considered significant.

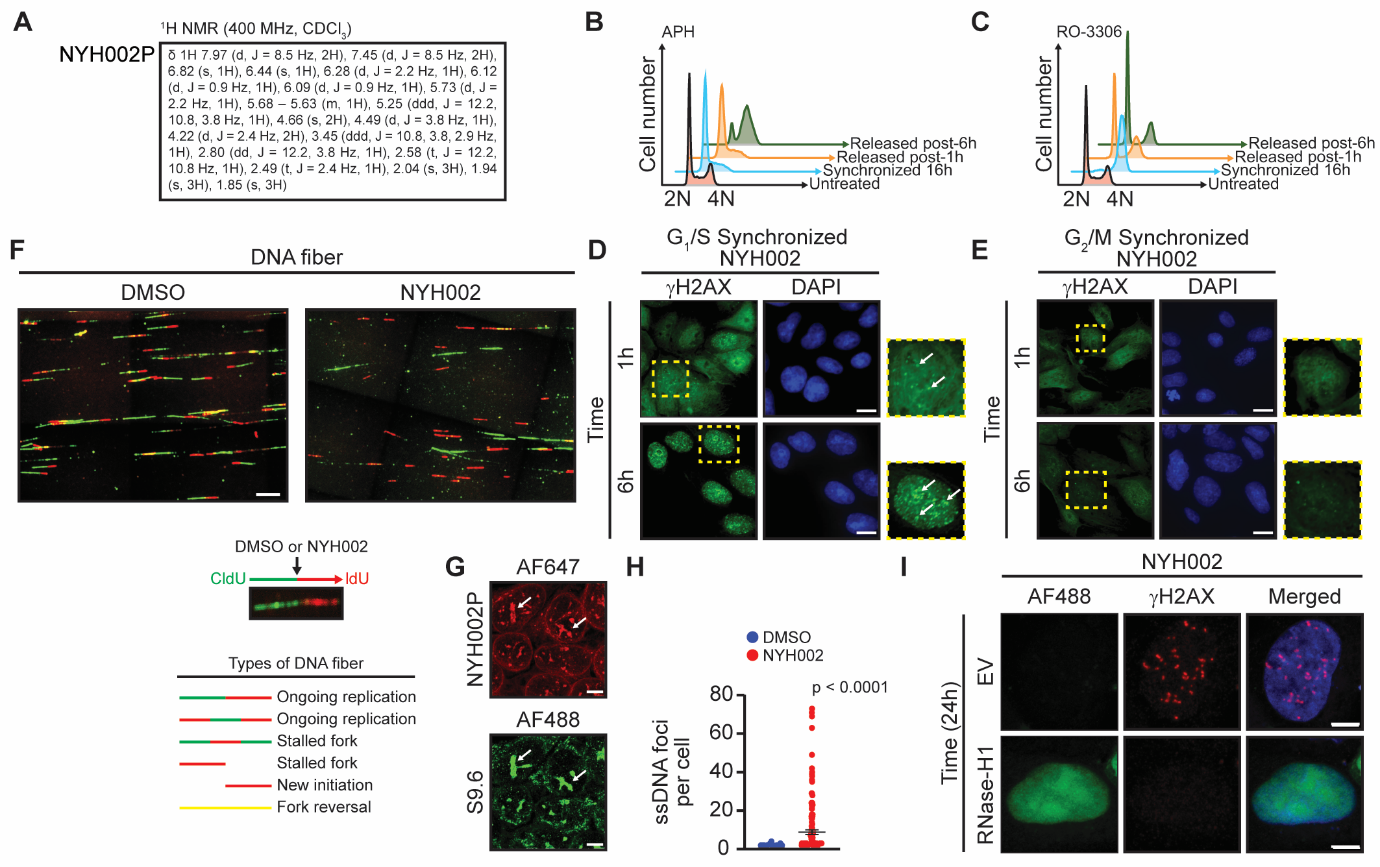

**Supplementary Figure 3:** Phenotypic profiling of replication stress.

(A) ^1^H NMR structural information of NYH002-alkyne probe (NYH002P).

(B, C) Synchronous cell cycle profile of propidium iodide-stained DLD1 cells at 16 hours with 10 µM APH to achieve G_1_/S-phase arrest or the use of 10 µM RO-3306 to arrest cells at G_2_-phase. Upon release (one and six hours), APH-treated cells entered S-phase, while RO-3306-treated cells entered G_1_-phase.

(D, E) Representative γH2AX immunofluorescence images of G_1_/S- or G_2_-phase synchronized DLD1 cells treated with 1 µM NYH002, as indicated by the different time points. Yellow dash boxes inform recruitment of γH2AX foci (arrows, white) to the site of DNA DSB. Scale bar: 10 μm.

(F) DNA fibers presented as immunofluorescence images of pulse-labeled DLD1 cells with CldU (green) and IdU (red), treated with either DMSO or 1 µM NYH002 for 30 mins. Scale bar: 20 μm.

(G) Fluorescence intensity of NYH200P-AF647 fluorophore and R-loop (arrows, white) as revealed by S9.6 staining at six hours post-treatment. Scale bar: 5 μm.

(H) Quantification of ssDNA foci per cell. DLD1 cells were pulse-labeled with 10 µM IdU for 24 hours, followed by treatment with 1 µM NYH002 for an additional 24 hours. DNA was processed under native conditions to detect ssDNA.

(I) Representative images of γH2AX-positive cells as a readout for replication-induced DSBs. DLD1 cells were transfected with either empty vector (EV) or RNaseH1 plasmid for 24 hours, followed by treatment with 1 µM NYH002. Scale bar: 5 μm.

Data are presented as mean ± SEM. Mann-Whitney U test (H). p-values < 0.05 were considered significant.

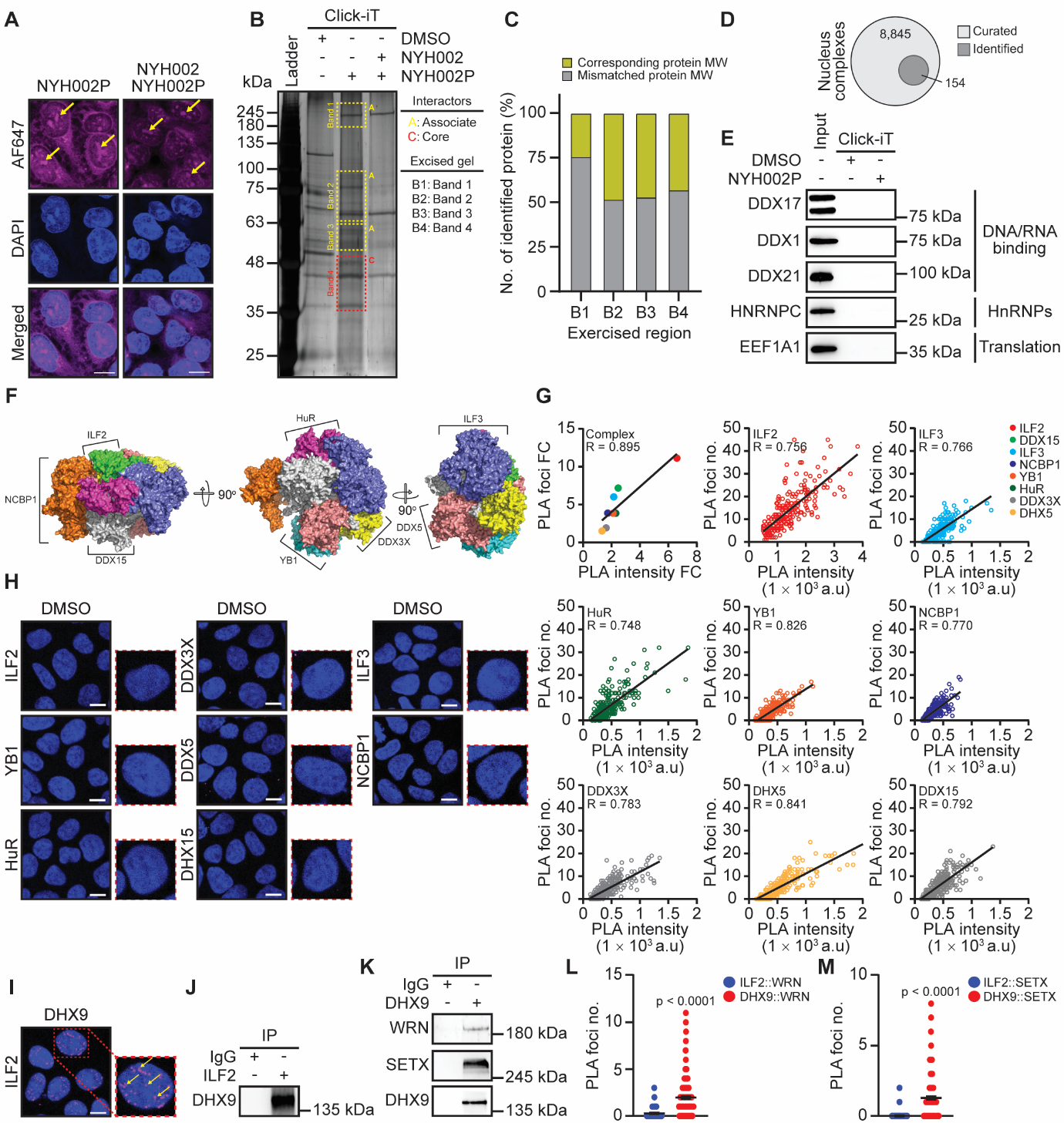

**Supplementary Figure 4:** NYH002 probe identifies ILF2–DHX9 axis.

(A) Representative immunofluorescence images of competitive binding between NYH002 and NYH002P-AF647 fluorescent probe (arrow, yellow) in the nucleus of DLD1 cells. Scale bar: 10 μm.

(B, C) A representative silver-stained gel of NYH002P co-immunoprecipitation with WCL of DLD1 cells. The red dash box indicates direct (core) drug–protein interactions, while the yellow dash boxes indicate indirect (associated) drug–protein interactors. All gel bands (B1–B4) were processed for label-free quantification and protein identification to match the corresponding molecular weight of the excised region.

(D) Interrogation of the CORUM database revealed that 154 out of 8,845 nuclear complexes contained at least one NYH002P-associated protein.

(E) Immunoblot of indirect candidate proteins through NYH002P immunoprecipitation with WCL of DLD1 cells. Proteins of interest were randomly selected from each functional group as shown. Input represents 1% of the total WCL.

(F) AlphaFold3 prediction of an *in silico* ILF2 complex.

(G) Determination of direct drug–protein interactions was based on the total number of PLA foci observed between NYH002P and the identified proteins of the ILF2 complex. A Pearson correlation coefficient greater than 0.7 indicates a high frequency of protein recruitment and activity (n = 3).

(H) Negative control for PLA between DMSO and the various proteins of the ILF2 complex.

(I) Representative image of PLA showing the interaction between ILF2 and DHX9 in DLD1 cells. Scale bar: 10 μm.

(J) Immunoprecipitation of ILF2 to detect DHX9 in WCL from DLD1 cells (n = 3).

(K) Co-immunoprecipitation of DHX9 to detect R-loop-associated helicases in WCL from DLD1 cells.

(L) Quantification of PLA foci comparing ILF2-WRN interactions to DHX9-WRN interactions (n = 3).

(M) Quantification of PLA foci comparing ILF2-SETX interactions to DHX9-SETX interactions (n = 3).

Data are presented as mean ± SEM. Mann-Whitney U test (L and M). p-values < 0.05 were considered significant.

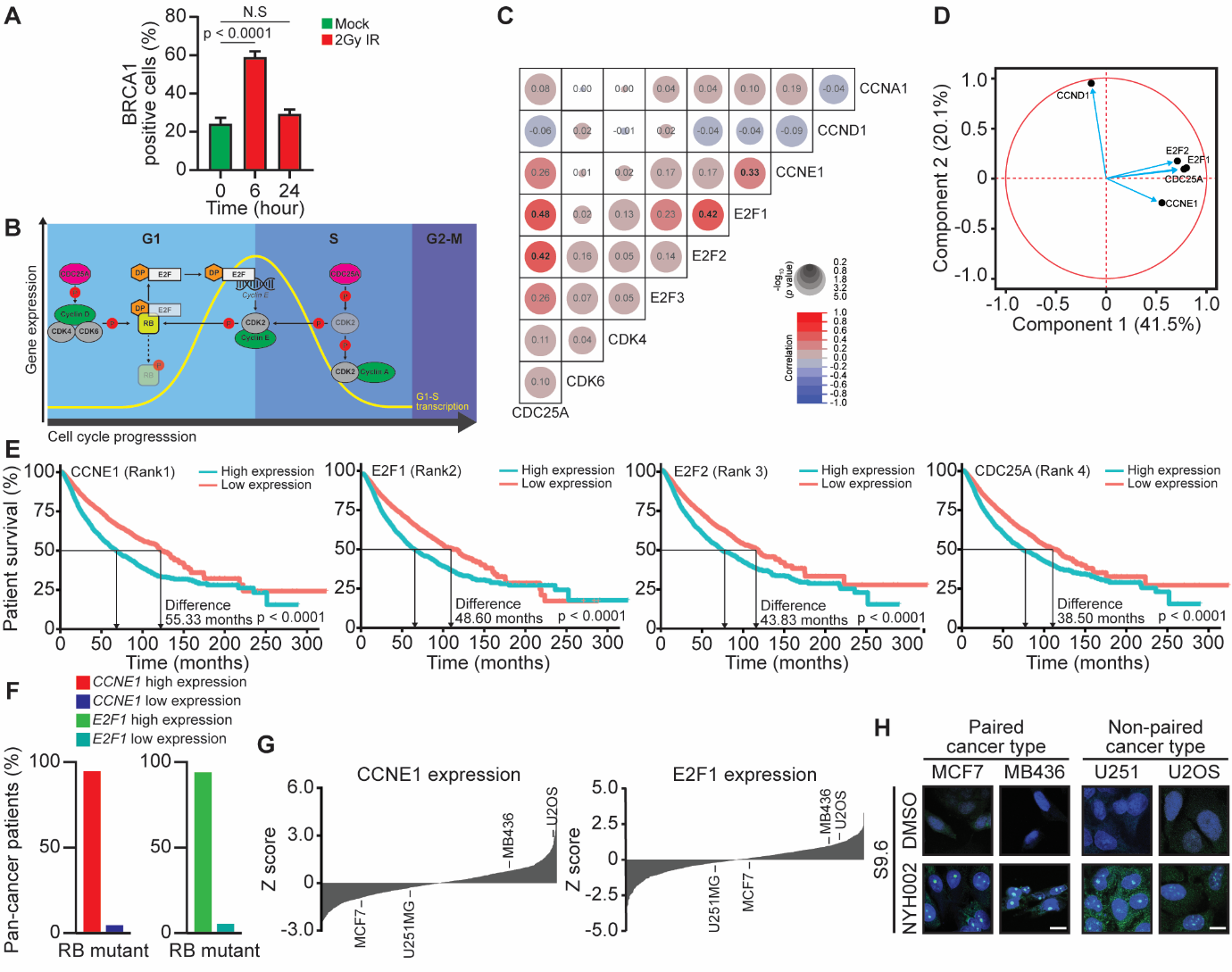

**Supplementary Figure 5:** Identification of G_1_/S checkpoint deficient cancer cells

(A) HR proficiency was determined by the induction and resolution of BRCA1-positive cells (> 5 foci per nucleus) following 2 Gy ionizing radiation (IR) in DLD1 cells.

(B) A simplified G_1_/S cell-cycle concept map.

(C) Gene expression correlation matrix for G_1_-, G_1_/S-, and S-phase cell cycle-related genes from the TCGA pan-cancer cohort. The size of the circle denotes significance, while the numerical readout informs the correlation value between genes.

(D) Principal component analysis plot of G_1_/S gene candidates with high correlation was based on the gene expression of the TCGA pan-cancer cohort. *CCND1* gene expression was used as a negative control.

(E) Kaplan-Meier (KM) plot of high and low expression levels of *CCNE1*, *E2F1*, *E2F2*, and *CDC25A*. Gene significance was ranked based on the greatest difference in 50% overall survival.

(F) *RB1-mutant* pan-cancer patients (%) from TCGA exhibit differences in *CCNE2* and *E2F1* gene expression.

(G) Selection of cancer cell lines with differential *CCNE1* and *E2F1* expressions (high vs low) was performed by Z-scoring cell lines from the Cancer Cell Line Encyclopedia (CCLE) database. A pair-matched cell line, MCF7 and MDA-MB-436, was selected to represent breast cancer, whereas a non-paired tumor comprising U251 and U2OS represents glioblastoma and osteosarcoma, respectively.

(H) Representative immunofluorescence images of S9.6 staining in cancer cells treated with either DMSO or 1 µM NYH002 for 24 hours. Scale bar: 10 μm.

Data are presented as mean ± SEM. Unpaired two-tailed t-test (A) and Mantel-Cox test (E). p-values < 0.05 were considered significant, whereas N.S was regarded as not significant.

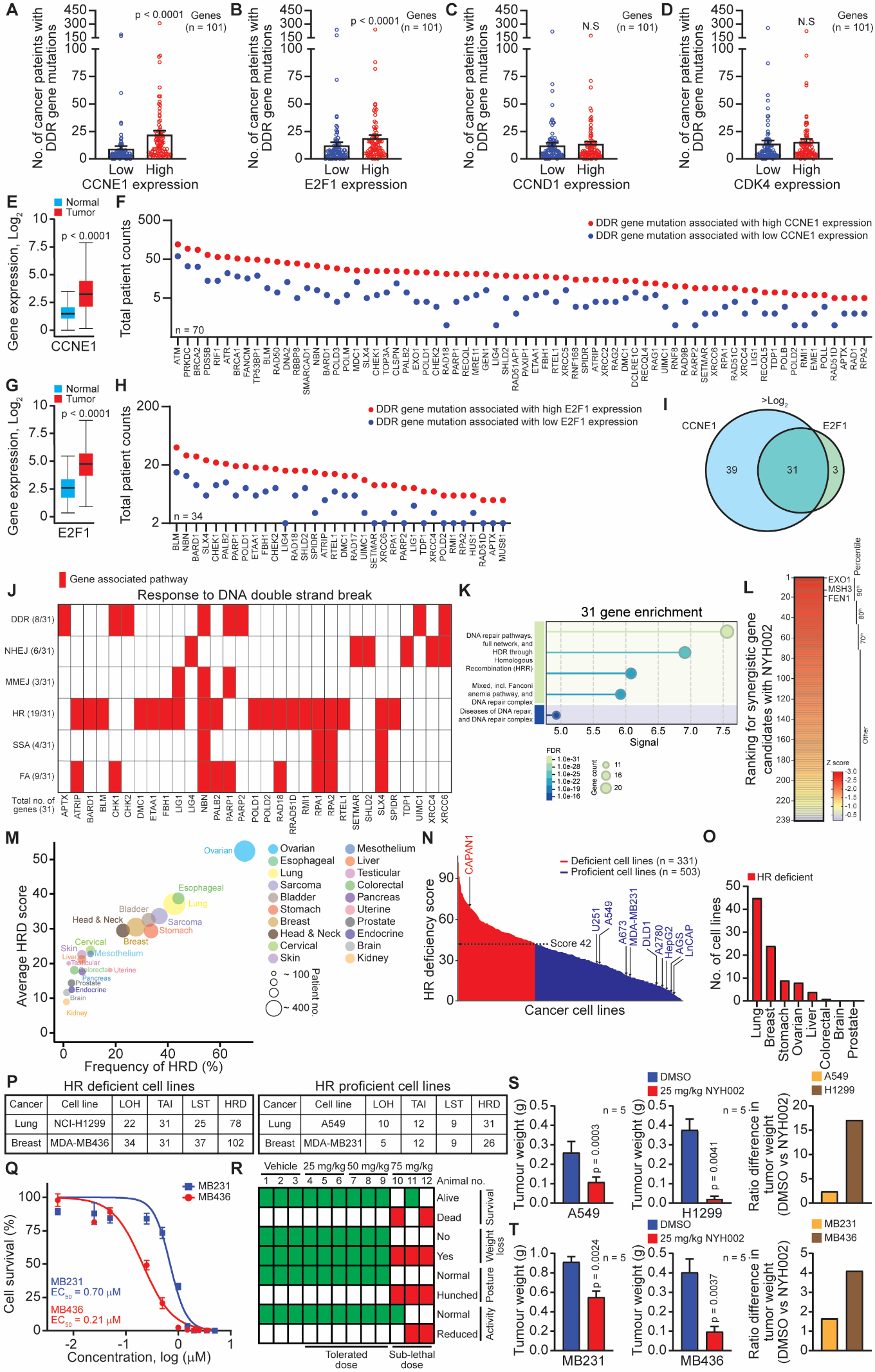

**Supplementary Figure 6:** Identification of HR proficient and deficient cell lines.

(A, B, C, D) Comparison of pan-cancer patients from TCGA exhibiting high or low expression of G_1_/S-phase genes, *CCNE2* and *E2F1*, in association with mutations in DDR genes. Expression of G_1_-phase genes, *CCND1* and *CDK4*, was used as a negative control to assess cell cycle phase specificity.

(E, F) Gene expression of *CCNE1* in normal and tumor samples from the TCGA cohort (left), including corresponding patients (right) with mutations in 70 out of 101 DDR genes.

(G, H) Gene expression of *E2F1* in normal and tumor samples from the TCGA cohort (left), including corresponding patients (right) with mutations in 34 out of 101 DDR genes.

(I) Venn diagram showing the overlap of mutated DDR genes (n = 31) in pan-cancer TCGA patients with high gene expression of *CCNE1* and *E2F1*.

(J, K) Mutated DDR genes associated with high expression of *CCNE1* and *E2F1*, and their involvement in various DNA DSB repair pathways, including pathway enrichment analysis using STRING.

(L) Z-score ranking of the DDR library (n = 239) that is based on the synergistic efficacy of individual siRNA genes and NYH002 treatment.

(M) An overall view of genomic scar score (loss of heterozygosity, LOH; telomeric allelic imbalance, TAI; large-scale transitions, LST) in different solid cancers from the TCGA cohort for HRD characterization.

(N) Ranking of HRD scores in the CCLE database and the selection of pan-cancer cell lines classified as either HR-proficient (< 42 score, in blue) or HRD (> 42 score, in red).

(O) Top eight most frequent number of HRD solid cancer cell lines in the CCLE database.

(P) Selection of HR-proficient and HRD pair-tumor cell lines was based on the genomic scar score (LOH, TAI, LST).

(Q) NYH002 dose-response curve of HR-proficient MDA-MB-231 and HRD MDA-MB-436 after seven days of treatment (n = 3).

(R) *In vivo* dose escalation in non-tumorigenic animals that received intraperitoneal injections on alternate days. Animal behaviour and weight were assessed to evaluate drug toxicity.

(S, T) *In vivo* therapeutic efficacy of NYH002 was assessed in HR-proficient and HRD tumor models for both lung and breast cancer. Individual tumor weights were recorded at the endpoint following a controlled cull.

Data are presented as mean ± SEM. Unpaired two-tailed t-test (A, B, C, D, E, G, S, and T). p-values < 0.05 were considered significant whereas N.S was regarded as not significant.
